# Metabolically Activated Proteostasis Regulators Reduce Differentiation of CD4^+^ T_H_17 Cells

**DOI:** 10.64898/2026.09.02.747634

**Authors:** Priyadarshini Chatterjee, Keishla Sánchez Ortiz, Stanislav Dikiy, Michael J. Bollong, Jessica E. Thaxton, Alejandra Mendoza, R. Luke Wiseman

## Abstract

The differentiation of naïve CD4^+^ T cells into effector T cell subsets such as T_H_1, T_H_2, T_H_17 and T_REG_ cells is governed by tightly coordinated activation programmes dependent on T cell receptor (TCR) engagement, co-stimulation, and cytokine signalling. This differentiation process involves regulation of stress-responsive signalling pathways such as the unfolded protein response (UPR) and the oxidative stress response (OSR) to adapt to the physiologic demands specific to distinct T cell subset functions. This suggests that pharmacologically targeting stress-responsive pathways offers a unique opportunity to selectively remodel the differentiation of these different T cell subsets. We previously identified compound AA147 as a metabolically activated proteostasis regulator that can induce both the ATF6 signalling arm of the UPR and, in certain cell types, the NRF2-regulated arm of the OSR.(Paxman et al., 2018; Plate et al., 2016) Here, we show that treatment with AA147 selectively reduces differentiation of pro-inflammatory T_H_17 cells by promoting degradation of the lineage-specifying transcription factor RORγt, without impacting the transcription factors of other CD4^+^ T cell subsets. We find that AA147-dependent reduction in T_H_17 differentiation is independent of ATF6 activation but involves NRF2 activation and reductions in intracellular reactive oxygen species (ROS). Apart from RORγt, we further show that AA147 decreases expression of the TCR-responsive factor IRF4, thus suppressing production of select effector cytokines across effector T cells, revealing additional ways in which this compound reshapes the activities of these essential T cell subsets. Our results demonstrate the potential for metabolically activated proteostasis regulators such as AA147 to selectively reshape T_H_17 cell identity while broadly dampening effector cytokine responses across effector T cell subsets.

## INTRODUCTION

CD4^+^ T cells are key players in adaptive immunity, as they not only coordinate immune effector functions across other immune cell populations, but also shape the nature and persistence of the immune response through their differentiation into specialised effector and memory states.(Zhu et al., 2010; Luckheeram et al., 2012; Schnell et al., 2023; Masopust et al., 2026) The differentiation of naïve CD4^+^ T cells into effector T cells is governed by tightly coordinated activation programmes dependent on T cell receptor (TCR) engagement, co-stimulation, and cytokine signalling, which gives rise to T_H_1, T_H_2, T_H_17 and T_REG_ cells.(Espinosa et al., 2020) This process requires rapid cellular reprogramming that drives expansion and differentiation, enabling effector functions such as migration to appropriate tissue sites and the production of signature cytokines such as IFNγ, IL-4, IL-17A, and IL-10.(Zhu et al., 2010; Saravia et al., 2019) While critical for their ability to drive immune responses against diverse pathologic insults, dysregulated T cell responses are implicated in the pathogenesis of many autoimmune and inflammatory diseases.(Chandwaskar et al., 2024; Farchione et al., 2026; Sligar et al., 2026) In particular, T_H_17 cells can be pathogenic in a variety of autoimmune and chronic inflammatory diseases such as psoriasis, inflammatory bowel disease (IBD) and multiple sclerosis (MS) which has led to considerable interest in identifying therapeutic opportunities to regulate their differentiation.(Louten et al., 2009; Vlachos et al., 2016; Fasching et al., 2017; Moser et al., 2020; Schnell et al., 2023)

T cell activation and differentiation impose significant amounts of endoplasmic reticulum (ER) and oxidative stress due to the increased production of secretory proteins and metabolic reprogramming associated with these processes.(Chen et al., 2023; Pino et al., 2008; Shu et al., 2023) To confront this, cells utilise stress-responsive signalling pathways such as the unfolded protein response (UPR) and oxidative stress response (OSR) to cope with the increased protein demand required for differentiation. The UPR comprises three signalling pathways regulated downstream of the ER membrane proteins IRE1, PERK, and ATF6.(Read and Schröder, 2021; Acosta-Alvear et al., 2025) In response to pathologic ER insults, these three UPR signalling pathways are activated to promote the adaptive remodelling of ER and cellular physiology primarily through the activation of their downstream transcription factors XBP1s, ATF4, and ATF6 (a cleaved product of full-length ATF6). The OSR is primarily regulated by the reactive oxygen species (ROS)-activated transcription factor NRF2.(Lo et al., 2006; Ma, 2013; Ngo and Duennwald, 2022; Zhang, 2025) In the absence of ROS, NRF2 is targeted to proteasomal degradation through the activity of the E3 ubiquitin ligase KEAP1. When ROS accumulates, KEAP1 is covalently modified by ROS on a regulatory cysteine, preventing the ubiquitination of NRF2 and allowing NRF2 to localise to the nucleus to induce expression of antioxidant genes.

The activity of UPR and OSR signalling pathways has been shown to reshape the differentiation and activity of T cell subsets in many different physiologic and disease contexts.(Kemp et al., 2013; Wu et al., 2023; Zhao et al., 2023; Wu et al., 2024.; Wu and Zhong, 2024; Cheng et al., 2025) For example, the unconventional activation of IRE1 through a JAK2-dependent mechanism promotes T_H_17 differentiation in the mouse models of airway epithelial inflammation.(Wu et al.,2024) IRE1 activity also promotes IL-4 expression in differentiating T_H_2 cells (Kemp et al., 2013), while ATF6 activity regulates T_H_2 and T_H_17 responses in mouse models of asthma.(Wu et al., 2023) Alternatively, in the context of graft versus host disease, PERK signalling contributes to the increased differentiation of T_H_1 and T_H_17 populations, and reduced differentiation of T_REG_ cells.(Cheng et al., 2025)

NRF2-dependent regulation of ROS is also critical for efficient differentiation of T_H_17 cells.(Zhao et al., 2023) The differentiation of T_H_17 cells requires a modest amount of ROS, while excessive ROS accumulation leads to cytotoxicity, highlighting the sensitivity of this T_H_ cell subset to NRF2-dependent regulation of cellular redox.(Wu and Zhong, 2024) Consistent with this, both reductions and increases in NRF2-dependent ROS regulation reduce differentiation of T_H_17 cells.(Zhao et al., 2016; Zhao et al., 2023) Apart from CD4^+^ T cells, UPR and OSR signalling are critical for regulating the differentiation and activity of other T cell subsets, including CD8^+^ T cells, highlighting the importance of these pathways for adapting these immune cell populations to their specific effector functions.(Shu et al., 2023; Zhang and Cao, 2025)

The sensitivity of specific T_H_ cell populations to different stress-responsive signalling pathways suggests that pharmacologically targeting these pathways offers a unique opportunity to reshape an immune response in response to a specific pathologic insult. We previously identified AA147 as a metabolically activated proteostasis regulator that could activate adaptive, protective signalling through the ATF6 arm of the UPR.(Plate et al., 2016) AA147 functions through a mechanism involving metabolic activation by ER oxidases to generate an electrophilic quinone methide at the ER membrane.(Kline et al., 2023; Paxman et al., 2018) This reactive electrophile covalently modifies a small set of protein disulfide isomerases (PDIs) responsible for regulating disulfides within the ATF6 luminal domain. AA147-dependent modification of these PDIs increases the population of monomeric ATF6 that can then traffic to the Golgi, where the active N-terminal transcription factor domain is proteolytically released by the site 1 and site 2 proteases.(Paxman et al., 2018; Kline et al., 2023) The ATF6 transcription factor domain then localises to the nucleus where it induces expression of target genes involved in many biological functions, most notably ER proteostasis regulation.(Adachi et al., 2008; Shoulders et al., 2013) While AA147 is highly selective for ATF6 activation in many cell lines and *in vivo* tissues, in certain cells, most notably neurons, metabolically activated AA147 can also covalently target KEAP1 to activate the NRF2-regulated OSR.(Rosarda et al., 2021; Kline et al., 2024) Thus, AA147 offers the unique opportunity to induce protective NRF2 and/or ATF6 signalling in different contexts.

AA147 has been widely used to probe the therapeutic benefits of increased ATF6 and/or NRF2 signalling in many cellular and *in vivo* models of different disorders. For example, AA147-dependent ATF6 activation promotes adaptive remodelling of ER proteostasis pathways to reduce the secretion and extracellular aggregation of destabilised, aggregation-prone variants of proteins including transthyretin (TTR) or α1-antitrypsin (A1AT).(Plate et al., 2016; Sun et al., 2023) ER proteostasis remodelling afforded by AA147-dependent ATF6 activation has also been shown to rescue the trafficking and plasma membrane activity of mutant GABA_A_ receptors involved in idiopathic epilepsy.(Wang et al., 2022) AA147-dependent ATF6 activation is also protective in retinal organoids models of eye diseases including achromatopsia and Macular Telangiectasia type 2 (MacTel).(Kroeger et al., 2021; Rosarda et al., 2023) Further, the activation of ATF6 by AA147 protects the heart from ischemia/reperfusion injury in mouse models.(Blackwood et al., 2019) In addition, AA147-dependent activation of both ATF6 and NRF2 integrates to promote adaptive remodelling of multiple cell types and tissues to protect in mouse models of diseases including Experimental Autoimmune Encephalomyelitis (EAE) and cerebral ischemia-reperfusion injury.(Yuan et al., 2022; Aksu et al., 2025) These results highlight the therapeutic potential for the metabolically activated proteostasis regulator AA147 to correct pathologic defects in multiple different cell types and tissues subjected to diverse types of pathologic insults.

Here, we sought to define the potential for AA147 to reshape CD4^+^ T cell differentiation through its activation of ATF6 and/or NRF2 signalling. We show that AA147 selectively suppresses the *ex vivo* differentiation of pro-inflammatory T_H_17 cells, without impacting the ability of T_H_1, T_H_2, or T_REG_ cells to acquire and maintain expression of lineage-specifying transcription factors. The AA147-dependent suppression of T_H_17 cells requires its metabolic activation but does not involve ATF6 activation in these differentiating cells. Instead, we find that AA147 primarily suppresses T_H_17 differentiation through a process involving the activation of NRF2. Mechanistically, we show that AA147 reduces intracellular ROS accumulation to promote proteasome-dependent degradation of the lineage defining T_H_17 transcription factor RORγt and in turn reduce expression of T_H_17 effector cytokines such as IL-17A. Additionally, we show that AA147 can reshape T cells responses more broadly by reducing the TCR-responsive transcription factor IRF4 and downstream effector cytokines in a NRF2 and ATF6 independent manner. Collectively, our results demonstrate that metabolically activated proteostasis regulators like AA147 can influence T cell responses by both reshaping T_H_17 differentiation and diminishing effector cytokine responses across effector T cell subsets.

## RESULTS

### AA147 reduces the ex vivo differentiation of CD4^+^ T_H_17 cells

We initially tested the impact of AA147 on the *ex vivo* differentiation of CD4^+^ T cells into different effector subsets. We isolated naïve CD4^+^ T cells from the spleens of C57BL/6J mice, activated them through TCR crosslinking using anti-CD3 and anti-CD28 antibodies, and then differentiated these cells in polarising conditions for T_H_1, T_H_2, T_H_17, and T_REG_ cell lineages using defined cytokine and antibody conditions in the presence or absence of AA147 (**Fig. 1A**).(Espinosa et al., 2020) T cell activation and proliferation depends on extracellular thiol availability and intracellular glutathione. Addition of antioxidants such as β-mercaptoethanol (BME) is essential to promote differentiation as it facilitates cysteine uptake and supports intracellular glutathione synthesis; however, exogenous addition of BME to cell culture media can suppress AA147 activity.(Angelini et al., 2002; Yan and Banerjee, 2010; Paxman et al., 2018) To accommodate this, we supplemented BME to differentiating cells 24 h after activation and addition of AA147, allowing enough time for this compound to become metabolically activated and covalently modify target proteins in differentiating CD4^+^ T cells (**Fig. 1A**). After 72 h of differentiation, we quantified the populations of cells expressing the lineage-specifying transcription factors T-bet (T_H_1), GATA-3 (T_H_2), RORγt (T_H_17), and Foxp3 (T_REG_) by flow cytometry. Treatment with AA147 did not influence the viability or activation of T cells toward these lineages (**Fig. S1A, B**). Further, AA147 did not influence expression of T-bet, GATA-3, or Foxp3 in cells differentiated in T_H_1, T_H_2, or T_REG_ cell polarising conditions, respectively (**Fig. S1C, D**). In contrast, treatment with AA147 reduced expression of the transcription factor RORγt in T cells polarised towards the T_H_17 cell lineage (**Fig. 1B, Fig. S1C**). AA147 similarly reduced production of the T_H_17 cytokines IL-17A and IL-22 in these cells (**Fig. 1C**, **Fig. S1E, F**). We performed Tandem Mass Tag (TMT)-based quantitative proteomics on CD4^+^ T cells polarised to T_H_17 cell lineages differentiated in the absence or presence of AA147 (**Table S1**). Gene ontology (GO) analysis showed that AA147 reduced the abundance of proteins involved in T_H_17 cell differentiation such as RORγt, IL-17A, and IL-17F, further confirming the AA147-dependent decrease in differentiation of this T cell subset (**Fig. 1D, E**).

**Figure 1.**
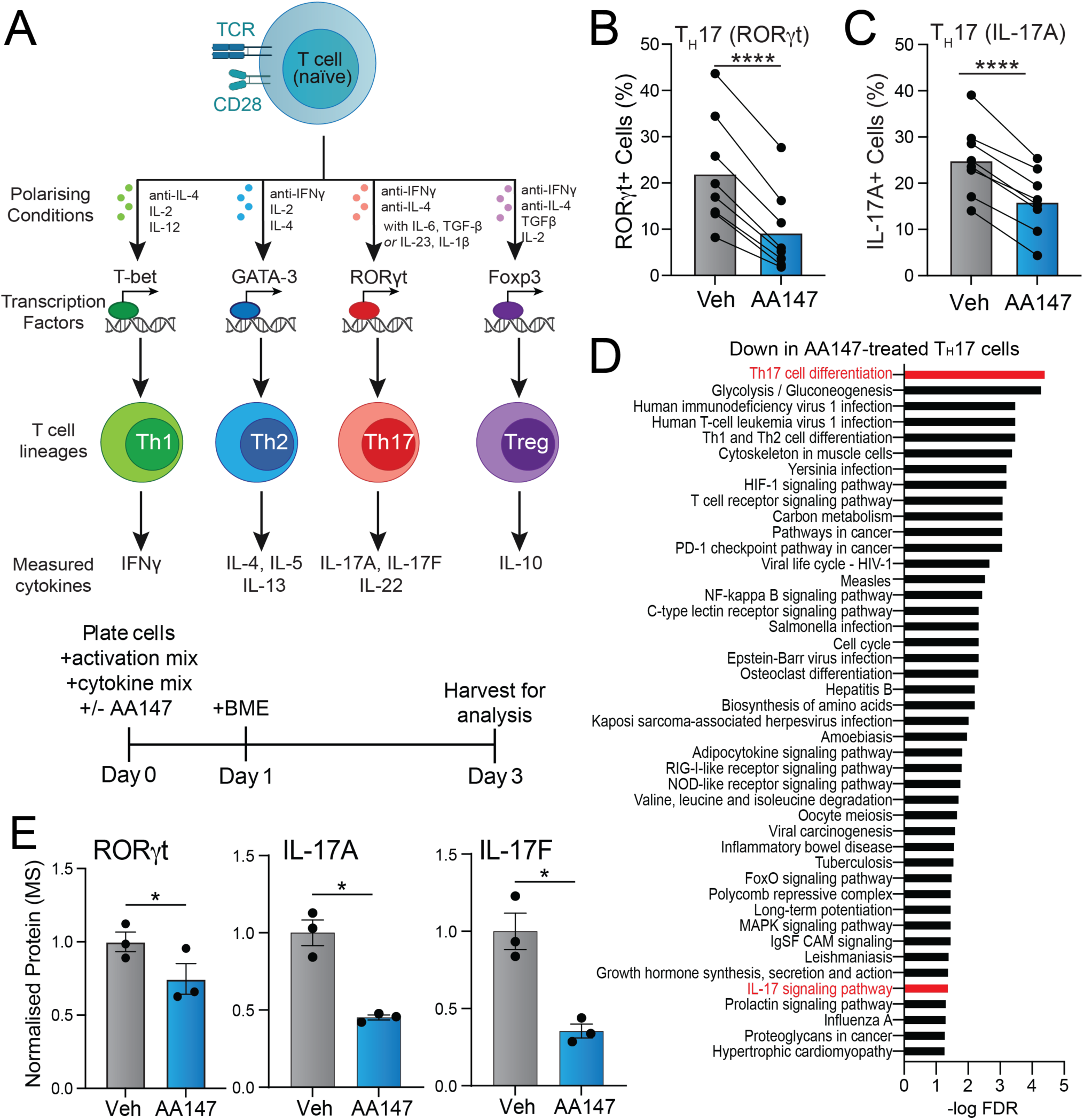
AA147 preferentially decreases differentiation of CD4^+^ T_H_17 cells. **A**. Schematic of CD4^+^ T cell lineage commitment. Naïve T cells differentiate into T_H_1, T_H_2, T_H_17 and T_REG_ cells upon treatment with specific polarising conditions. These cells are characterised by expression of their lineage defining transcription factors, T-bet, GATA-3, RORγt and Foxp3, respectively, and the production of specific effector cytokines. The timeline of our differentiation protocol is shown below. **B.** Percentage of RORγt^+^ T_H_17 cells differentiated in the presence of vehicle or AA147 (30 µM) for 72 h. Data are shown for CD4^+^ naïve T cells isolated from n=8 mice. **C.** Percentage of IL-17A^+^ T_H_17 cells differentiated in the presence of vehicle or AA147 (30 µM) for 72 h. Data are shown for CD4^+^ naïve T cells isolated from n=8 mice. **D**. KEGG pathway enrichment analysis of TMT-MS proteomic data identifying specific pathways downregulated in T_H_17 cells differentiated in the presence of AA147 relative to vehicle, ranked by -log(FDR). **E**. Relative protein levels, measured by TMT-MS, of RORγt, IL-17A, or IL-17F in T_H_17 cells differentiated in the presence of vehicle or AA147 (30 µM). Data are shown for CD4^+^ naïve T cells isolated from n=3 mice. *p<0.05, ***p<0.001, ****p<0.0001 for paired t-tests. Data described in panels B, D, and E are provided in **Source Data** Figure 1.

T_H_17 differentiation can be induced by the addition of multiple different cytokine combinations including IL-6/TGF-β or IL-23/IL-1β (**Fig. 1A**).(Espinosa et al., 2020) AA147 reduced expression of RORγt and IL-17A in T_H_17 T cells using either of these combinations of polarising cytokines (**Fig. S1G).** In contrast, AA147 did not influence expression of RORγt in already differentiated T_H_17 cells treated for 3 days, indicating that AA147 specifically disrupts the differentiation process but not the maintenance of the T_H_17 programme (**Fig. S1H**). Collectively, these results show that AA147 treatment reduces the *ex vivo* differentiation of T_H_17 cells relative to other T cell subsets.

### AA147 promotes proteasomal degradation of RORγt

RORγt protein levels can be regulated at the transcriptional and posttranslational levels.(Ciofani et al., 2012; Rutz et al., 2016) *Rorc*, the gene that encodes RORγt, is transcriptionally induced during T_H_17 cell differentiation by the activity of the transcription factor phosphorylated STAT3 (pSTAT3).(Durant et al., 2010) We found that AA147 treatment does not influence pSTAT3 levels in differentiating T_H_17 cells treated with IL-6 (**Fig. 2A,B**). Furthermore, AA147 did not reduce the expression of *Rorc* or decrease the expression of the alternative pSTAT3 target gene *Maf* in differentiating T_H_17 cells (**Fig. 2C, Fig S2A**). These results suggest that AA147 does not reduce RORγt expression by decreasing its transcription downstream of pSTAT3. However, AA147 does reduce the expression of the RORγt target gene *IL17a*, reflecting its ability to influence RORγt stability and activity (**Fig. 2C**).

**Figure 2.**
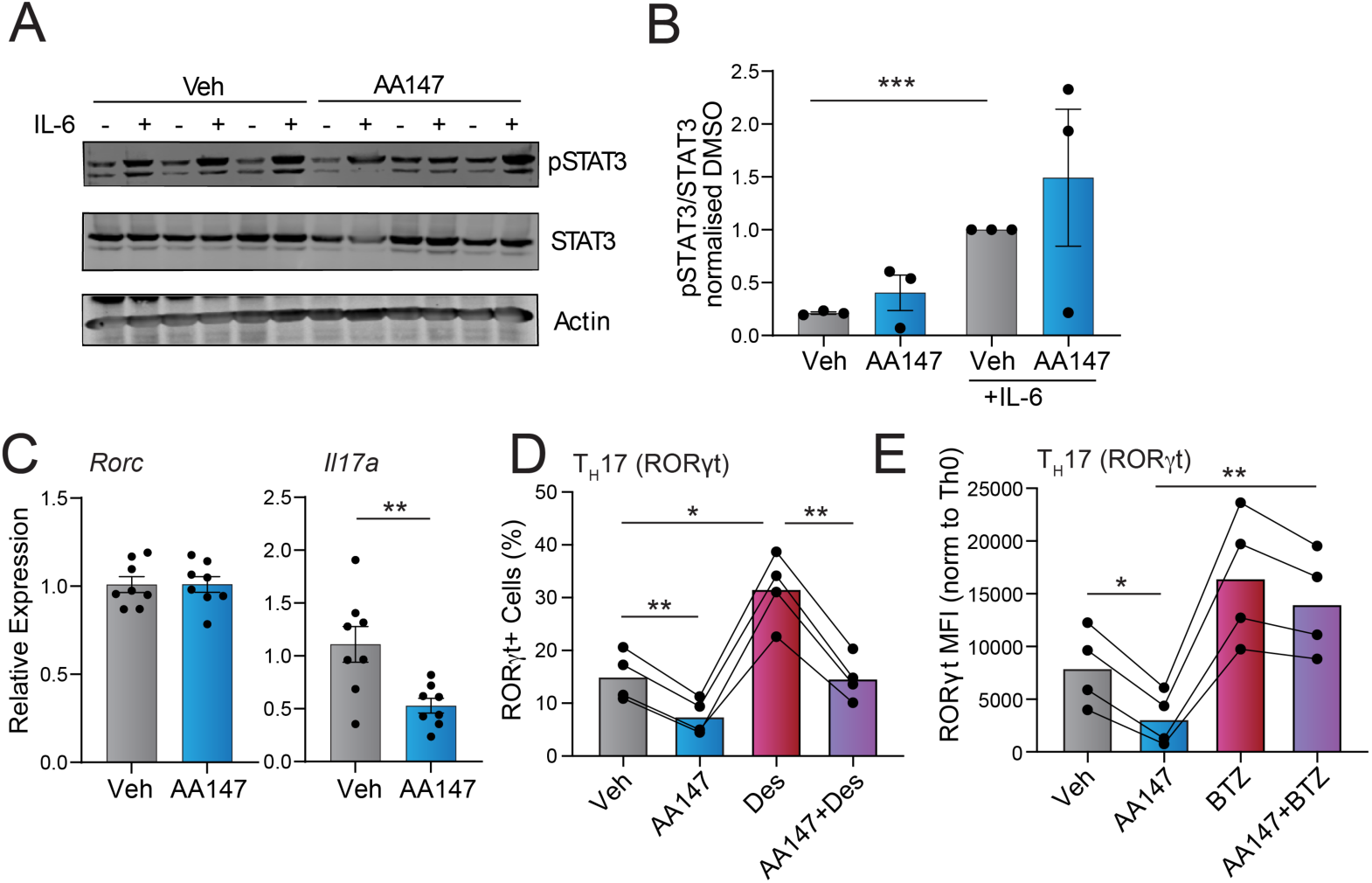
AA147 promotes proteasomal degradation of RORγt. **A, B.** Representative immunoblot (**A**) and quantification (**B**) of pSTAT3 in lysates prepared from T_H_17 cells stimulated for 15 minutes at 37°C with IL-6 (20 ng/mL) in the absence or presence of AA147 (30 µM). Data are shown for CD4^+^ T cells isolated from n=3 mice. **C.** Expression, measured by RT-qPCR, of *Rorc* or *IL17a* in CD4^+^ T cells differentiated for 72 h in T_H_17 polarising conditions in the presence or absence of AA147 (30 µM). Data for CD4^+^ T cells isolated from n=8 mice are shown. **D**. Percentage of CD4^+^ RORgt^+^ T_H_17 cells differentiated for 72 h in the presence of AA147 (30 µM) and/or desmosterol (10 µM), as indicated. Data are shown for CD4^+^ T cells isolated from n=4 different mice. **E**. Mean fluorescent intensity (MFI) for RORγt in T_H_17 cells differentiated for 72 h in the absence or presence of AA147 (30 µM) and then treated for 4 h with the proteasome inhibitor bortezomib (BTZ; 10 µM). The MFI was normalised to that observed in activated T_H_0 CD4^+^ T cells. Data are shown for CD4^+^ naïve T cells isolated from n=4 different mice. *p<0.05, **p<0.01, ***p<0.005, ****p<0.005 for one-way ANOVA (panels **B**, **D, E**) or paired t-test (panels **C**). Data described in panels B-E are provided in **Source Data** Figure 2.

RORγt is a member of the nuclear receptor family of transcription factors that is stabilised and activated by endogenous ligands such as the cholesterol precursor desmosterol.(Hu et al., 2015). We tested if AA147-dependent reductions in RORγt could be attributed to its impaired stabilisation by monitoring the impact of co-treatment with desmosterol on T_H_17 cell differentiation. As expected, treatment with desmosterol on its own increased RORγt levels in differentiating T_H_17 cells, reflecting the predicted increase in stability of this nuclear receptor afforded by this treatment (**Fig. 2D**). Despite this increase, AA147 still reduced RORγt protein levels during co-treatment with desmosterol, indicating that the AA147-dependent reduction in RORγt is independent of changes to its binding of stabilising ligands (**Fig. 2D**).

We next tested whether AA147-dependent reductions in RORγt levels are mediated by increasing its targeting for proteasomal degradation. T_H_17 cells differentiated for 72 h and then treated for 4 h with the proteasome inhibitor bortezomib (BTZ) showed increased basal protein levels of RORγt (**Fig. 2E**). This corresponded with a modest increase in expression of *Rorc* (**Fig. S2B**). Intriguingly, T_H_17 cells differentiated in the presence of AA147 and then treated for 4 h with BTZ did not show reductions in RORγt protein (**Fig. 2E**). This is despite the relative expression of *Rorc* mRNA being similar in BTZ-treated T_H_17 cells differentiated in the presence or absence of AA147 (**Fig. S2B**). These results show that AA147-dependent reductions in RORγt can be inhibited by short treatments with the proteasome inhibitor BTZ, indicating that this decrease in RORγt protein can be attributed to increased targeting to proteasomal degradation.

### AA147-dependent reductions in T_H_17 cell differentiation are independent of ATF6 activation

AA147 activates the ATF6 arm of the UPR through its metabolic activation and covalent modification of ER-localised PDIs.(Paxman et al., 2018; Kline et al., 2023) We asked if AA147 reduced differentiation of T_H_17 cells through this same mechanism. We used an alkyne containing AA147 analogue (AA147^yne^; **Fig. S3A**) to probe the metabolic activation and covalent protein targeting of AA147 after its metabolic activation in differentiating T_H_17 cells.(Paxman et al., 2018) AA147^yne^ reduced IL-17A levels in differentiating T_H_17 cells, confirming that this analogue retains the ability to suppress T_H_17 cell differentiation (**Fig. S3B**). We treated differentiating T_H_17 cells with AA147^yne^ for 24 h and then monitored covalent protein labelling by appending a fluorophore to the alkyne moiety of modified proteins using click chemistry, as previously described.(Paxman et al., 2018; Kline et al., 2023) This showed that AA147^yne^ covalently modified proteins in differentiating T_H_17 cells (**Fig. 3A**). Moreover, co-treatment with AA147 reduced AA147^yne^ covalent protein modifications, demonstrating that these compounds target overlapping proteins in these cells (**Fig. 3A**). Consistently, co-treatment with BME, which blocks AA147^yne^-dependent protein labelling (Paxman et al., 2018; Kline et al., 2023), reduced the population of AA147^yne^ modified proteins in differentiating T_H_17 cells (**Fig. 3B**). These results indicate that AA147 can undergo metabolic activation and covalent protein modification in this CD4^+^ T cell population.

**Figure 3.**
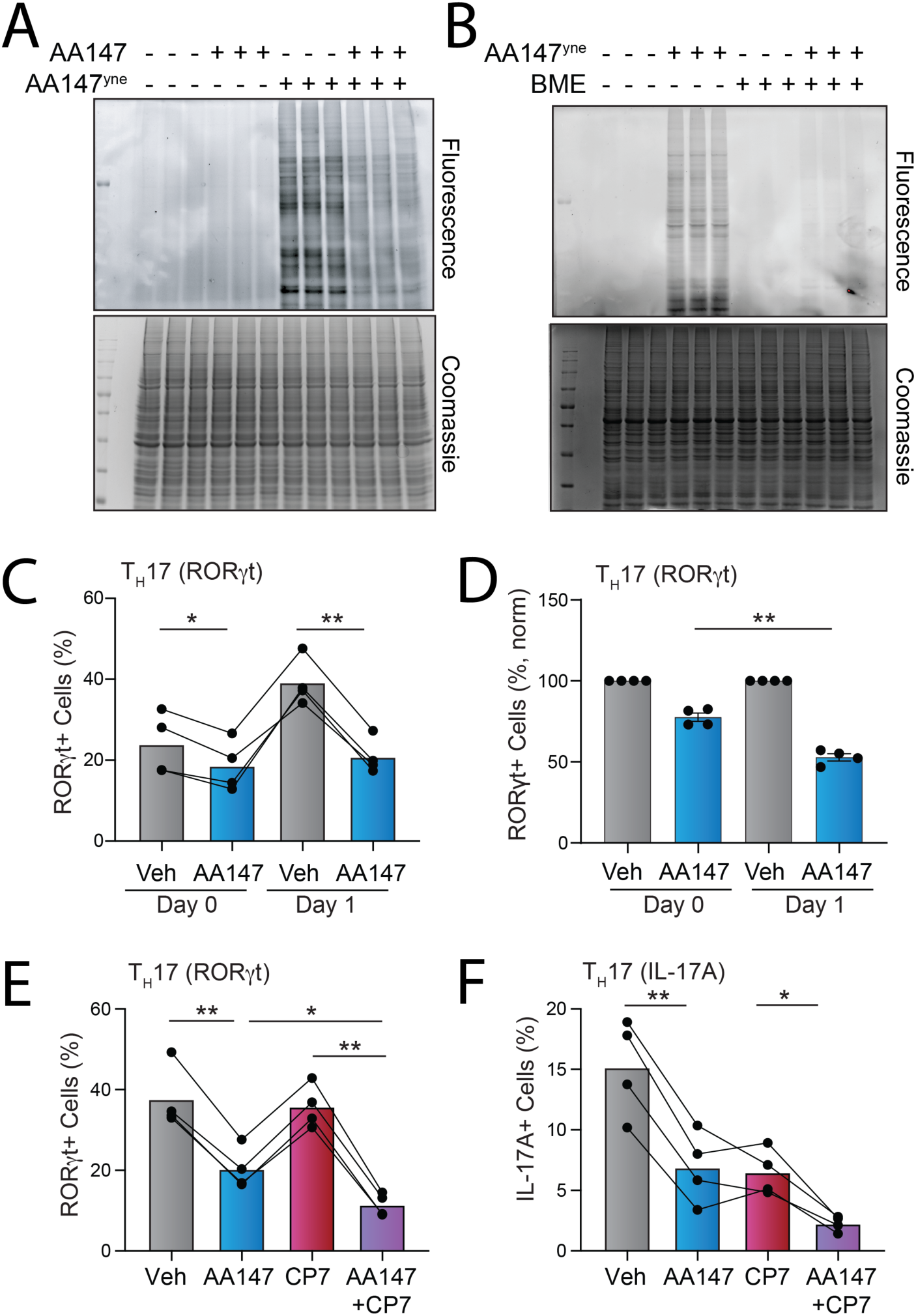
AA147-dependent reductions in T_H_17 differentiation are independent of ATF6 activation. **A.** Fluorescence and Coomassie gel of lysates prepared from naïve CD4^+^ T cells differentiated for 6 h in T_H_17 polarising conditions in the absence or presence of AA147 (30 µM) and then treated an additional 18 h with AA147^yne^ (10 µM), as indicated. **B**. Fluorescence and Coomassie images of gels prepared from naïve CD4^+^ T cells differentiated for 24 h in T_H_17 polarising conditions in the presence of AA147^yne^ (30 µM) and/or BME (20 µM), as indicated. **C, D**. Raw (left) and normalised percentage of CD4^+^ RORgt^+^ T_H_17 cells differentiated for 72 h in the presence of AA147 (30 µM) where BME was added either at Day 0 or Day 1, as indicated. Data are shown for CD4^+^ T cells isolated from n=4 different mice. **E, F**. Percentage of CD4^+^ RORγt^+^ (**E**) or IL-17A+ (**F**) T_H_17 cells differentiated for 72 h in the presence of AA147 (30 µM) and/or Ceapin-A7 (CP7, 3 µM), as indicated. Data are shown for CD4^+^ T cells isolated from n=4 different mice. *p<0.05, **p<0.01, ***p<0.005, ****p<0.005 for one-way ANOVA (panels **C, E, F**) or paired t-test (panel **D**). Data described in panels C-F are provided in **Source Data** Figure 3.

To test whether metabolic activation and covalent protein targeting by AA147 functionally contributed to the reduction of T_H_17 cell differentiation mediated by this compound, we compared AA147-dependent reductions in differentiation of T_H_17 cells incubated with BME from the beginning of the differentiation process (Day 0), where AA147-dependent protein labelling is inhibited (**Fig. 3B**), to that observed when BME was added after the start of differentiation (Day 1). We found that co-treatment with BME at Day 0 attenuated the AA147-dependent decrease in T_H_17 cell differentiation (**Fig. 3C, D**). This suggests that the reduction in T_H_17 differentiation afforded by AA147 is mediated through a mechanism involving its metabolic activation and covalent protein modification.

Next, we sought to determine the dependence of AA147-mediated reductions of T_H_17 cell differentiation on ATF6 activation using the selective ATF6 inhibitor Ceapin-A7 (CP7).(Gallagher et al., 2016.; Torres et al., 2019) We confirmed the increased expression of the ATF6 target gene *Hspa5* in T_H_17 cells differentiated for 3 days in the presence of AA147 (**Fig. S3C**). Co-treatment with CP7 blocked this AA147-dependent increase in *Hspa5*, demonstrating the effectiveness of this compound for blocking ATF6 activation during *ex vivo* T cell differentiation. Treatment with CP7 in the absence or presence of AA147 did not influence the viability or activation of differentiating T_H_17 cells (**Fig. S3D, E**). Treatment with CP7 alone did not influence RORγt expression in these cells (**Fig. 3E**). However, co-treatment with CP7 enhanced AA147’s suppression of RORγt expression in differentiating T_H_17 cells (**Fig. 3E**). Further, we found that treatment with CP7 basally reduced IL-17A expression in differentiating T_H_17 cells (**Fig. 3F**). This is consistent with previous results showing that CP7 decreases IL-17A expression in T_H_17 cells.(Wu et al., 2023; Yan et al., 2024). However, co-treatment with CP7 and AA147 further reduced IL-17A expressing to levels greater than that observed with either treatment alone (**Fig. 3F**). These results indicate that the reductions in T_H_17 differentiation afforded by AA147 are unlikely to be attributed to ATF6 activation.

### AA147 activates the NRF2 oxidative stress response to suppress T_H_17 differentiation

To gain insight into how AA147 impacts T_H_17 cell differentiation, we performed RNA sequencing (RNAseq) on T_H_17 cells differentiated 72 h after treatment with AA147 or vehicle (**Table S2**). As expected, AA147 reduced expression of RORγt target genes (e.g., *Il17a*, *Il17f*) (**Fig. 4A**, **Table S3**). We did not observe global increases in the expression of IRE1/XBP1s or ATF6 target gene sets (**Fig. 4B, Table S3**).(Grandjean et al., 2019) Further, AA147 did not influence expression of target gene sets regulated by other stress pathways such as the heat shock response (HSR) or the hypoxic stress response. We did observe increased expression of target genes regulated by the integrated stress response (ISR) and the OSR in T_H_17 cells (**Fig. 4B**, **Table S3**). Thus, we next sought to define the specific contributions of these two stress-responsive signalling pathways on AA147-dependent reductions in T_H_17 cell differentiation.

**Figure 4.**
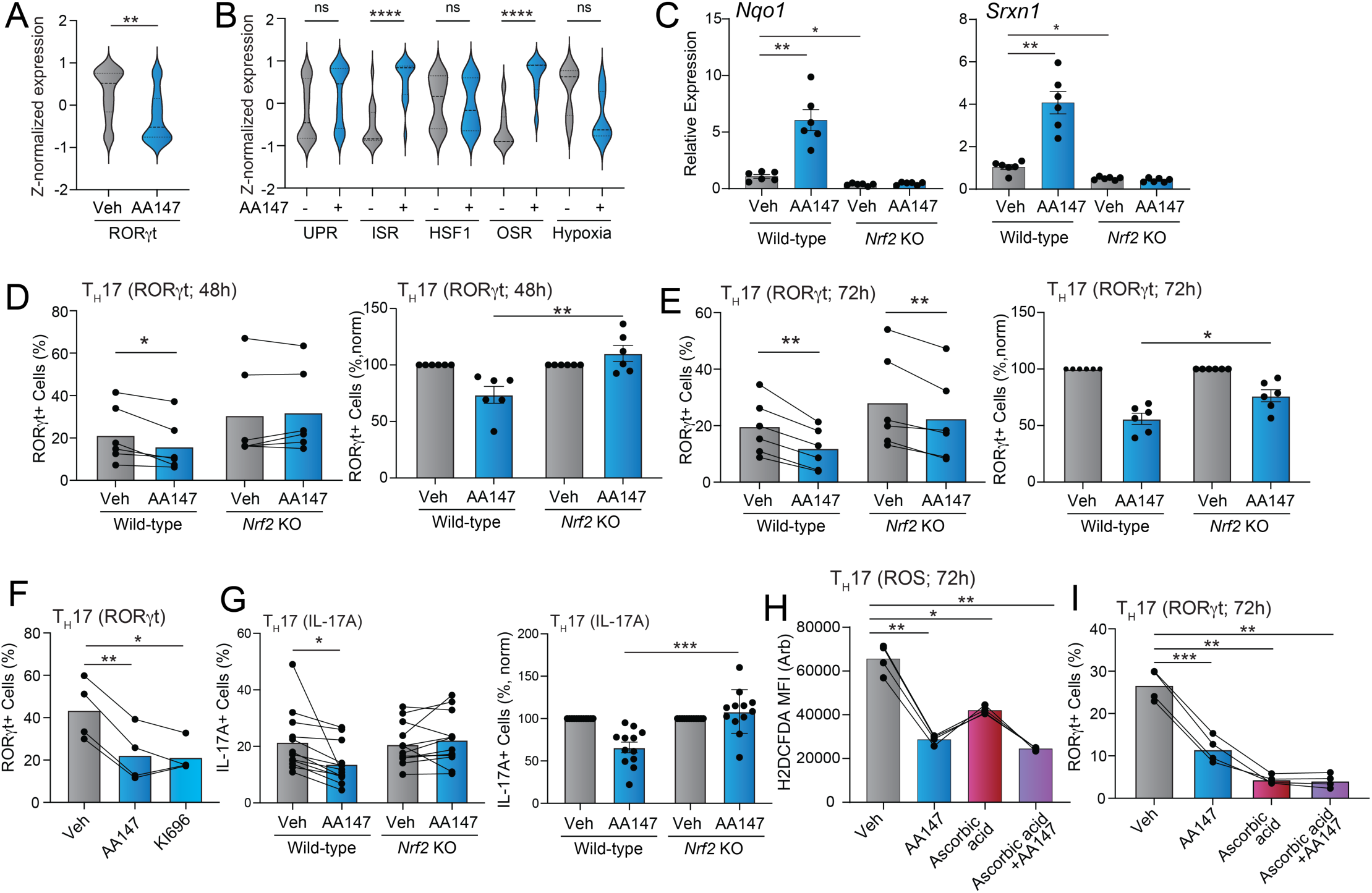
AA147 reduces T_H_17 differentiation through an NRF2-dependent mechanism. **A.** Z-normalised expression, measured by RNAseq, of RORγt target genes in T_H_17 cells differentiated in the presence or absence of AA147 for 72 h. A full list of RORγt target genes is shown in **Table S3**. **B**. Z-normalised expression, measured by RNAseq, of a geneset comprising target genes regulated by the ATF6 and IRE1/XBP1s arms of the UPR, the ISR, the HSR, the OSR, or the hypoxia stress response in T_H_17 cells differentiated in the presence or absence of AA147 for 72 h. A full list of target genes for these stress pathways is shown in **Table S3**. **C**. Expression, measured by RT-qPCR, of the NRF2 target genes *Nqo1* and *Srxn1* in CD4^+^ T cells isolated from wild-type or *Nrf2* KO mice differentiated for 72 h in T_H_17 polarising conditions in the presence or absence of AA147 (30 µM). Data for CD4^+^ T cells isolated from n=6 mice are shown. **D, E**. Raw (left) and normalised (right) percentage of CD4^+^ RORgt^+^ T_H_17 cells differentiated from wild-type or *Nrf2* KO mice for 48 h (**D**) or 72 h (**E**) in the presence of AA147 (30 µM). Data are shown for CD4+ T cells isolated from n=6 mice. **F**. Percentage of CD4^+^ RORgt^+^ T_H_17 cells differentiated for 72 h in the presence of AA147 (30 µM) or the NRF2 activator KI696 (5 µM), as indicated. Data are shown for CD4+ T cells isolated from n=4 different mice. **G.** Raw (left) and normalised (right) percentage of CD4^+^ IL-17A^+^ in T_H_17 cells differentiated from wild-type or *Nrf2* KO mice for 72 h in the presence of AA147 (30 µM). Data are shown for CD4^+^ T cells isolated from n=12 mice. **H.** Mean fluorescent intensity (MFI) for H_2_DCFDA staining in T_H_17 cells differentiated for 72 h in the absence or presence of AA147 (30 µM) and/or ascorbic acid (250 µM), as indicated. The MFI was normalised to that observed in activated T_H_0 CD4^+^ T cells. Data are shown for CD4^+^ T cells isolated from n=4 mice. **I.** Percentage of CD4^+^ RORgt^+^ T_H_17 cells differentiated for 72 h in the presence of AA147 (30 µM) and/or ascorbic acid (250 µM), as indicated. Data are shown for CD4^+^ T cells isolated from n=4 different mice. *p<0.05, **p<0.01, ***p<0.005, ****p<0.005 for one-way ANOVA (panels **B**, **C**, **F, H, J**) or paired t-test (panel **A** and raw data in panels **D, E, G**) or unpaired t-test (normalised data in panels **D, E**). Data described in panels C-I are provided in **Source Data** Figure 4.

ISR activation has been previously suggested to suppress T_H_17 cell differentiation.(Sundrud et al., 2009; Carlson et al., 2014) We confirmed that AA147 increased expression of the ISR target genes *Asns*, *Chac1*, and *Chop/Ddit3* in differentiating T_H_17 cells (**Fig. S4A**). Co-treatment with the ISR inhibitor ISRIB reduced AA147-dependent activation of these genes, demonstrating that their ISR-specific activation by AA147 (**Fig. S4A**). (Sidrauski et al., 2013; Sekine et al., 2015; Sidrauski et al., 2015) Although treatment with ISRIB basally increased T_H_17 cell differentiation, co-treatment with ISRIB did not influence AA147-dependent reductions in the differentiation of this T_H_ cell subset (**Fig. S4B**), suggesting that AA147 is unlikely to reduce T_H_17 cell differentiation through a mechanism involving increased ISR signalling.

The data motivated us to define the dependence of AA147-mediated reductions in T_H_17 cell differentiation on NRF2 (*Nfe2l*), the main transcriptional mediator of the OSR. We differentiated CD4^+^ T cells isolated from NRF2-deficient mice to T_H_17 cells in the absence or presence of AA147 for 48 or 72 h. We confirmed that AA147-dependent induction of NRF2 target genes *Nqo1* and *Srxn1* was blunted in NRF2-deficient T_H_17 cells (**Fig. 4C**). Treatment with AA147 did not influence viability or activation of NRF2-deficient T_H_17 cells (**Fig. S4C, D**). However, NRF2 deficiency attenuated the AA147-dependent reduction in T_H_17 cell differentiation at both 48 and 72 h (**Fig. 4D, E**). This indicates that AA147 reduces T_H_17 cell differentiation through an NRF2-dependent mechanism. Consistent with this, treatment with the alternative NRF2 activator KI696 which acts by competitively occupying the NRF2 binding pocket on KEAP1, similarly reduced T_H_17 cell differentiation (**Fig. 4F**).(Yasuda et al., 2025) NRF2 deficiency rescued IL-17A expression in differentiating T cells treated with AA147, mimicking the rescue of RORγt expression and activity, suggesting that the reductions in RORγt and IL-17A are both dependent on NRF2 (**Fig. 4G**)

NRF2 regulates the expression of antioxidant genes that function to reduce ROS.(Lo et al., 2006; Ma, 2013; Ngo and Duennwald, 2022; Zhang, 2025). Thus, we predicted that AA147-dependent reductions in T_H_17 cell differentiation could be attributed to decreased ROS. Consistent with this, AA147 treatment reduced ROS in differentiating T_H_17 cells (**Fig. 4H, Fig. S4E**). Treatment with the antioxidant ascorbic acid 2-phosphate also reduced ROS and suppressed RORγt expression in differentiating T_H_17 cells (**Fig. 4H, I, Fig. S4E**). However, co-treatment with AA147 and ascorbic acid 2-phosphate did not show further reductions in ROS or T_H_17 differentiation greater than that observed with ascorbic acid 2-phosphate alone, which suggests that these two treatments suppress T_H_17 differentiation through similar mechanisms (**Fig. 4H, I, Fig. S4E**). Collectively, these data align with a known role of ROS in regulating T_H_17 differentiation (Zhao et al., 2023; Wu and Zhong, 2024) and are consistent with a model whereby AA147 suppresses T_H_17 differentiation through a mechanism involving reductions of ROS.

### AA147 reduces IRF4 in CD4^+^ T cells through an NRF2-independent mechanism

Unlike what was observed for IL-17A, we found that mRNA and protein levels of IL-22 were decreased in both WT and NRF2-deficient T_H_17 cells treated with AA147 (**Fig. 5A, B**). This suggests that apart from NRF2 activation, AA147 could also influence T_H_17 cell effector functions through other mechanisms. Apart from RORγt, multiple other transcription factors, including BATF, AHR, and IRF4 are involved in the differentiation of T_H_17 cells.(Ciofani et al., 2012) We sought to define the impact of AA147 on the intracellular levels of these other T_H_17 transcriptional regulators. AA147 did not influence intracellular levels of AHR or BATF (**Fig. S5A, B**). However, we found that IRF4 levels were decreased in AA147-treated T_H_17 cells (**Fig. 5C, Fig. S5C**). Co-treatment with the ATF6 inhibitor Ceapin-A7 did not influence AA147-dependent decreases of IRF4 in T_H_17 cells (**Fig. 5C**). Further, we found that NRF2 deficiency also did not influence AA147-dependent reductions in IRF4 in differentiating T_H_17 cells (**Fig. 5D**). These results indicate that AA147 decreases IRF4 expression through a mechanism independent of ATF6 or NRF2 activation.

**Figure 5.**
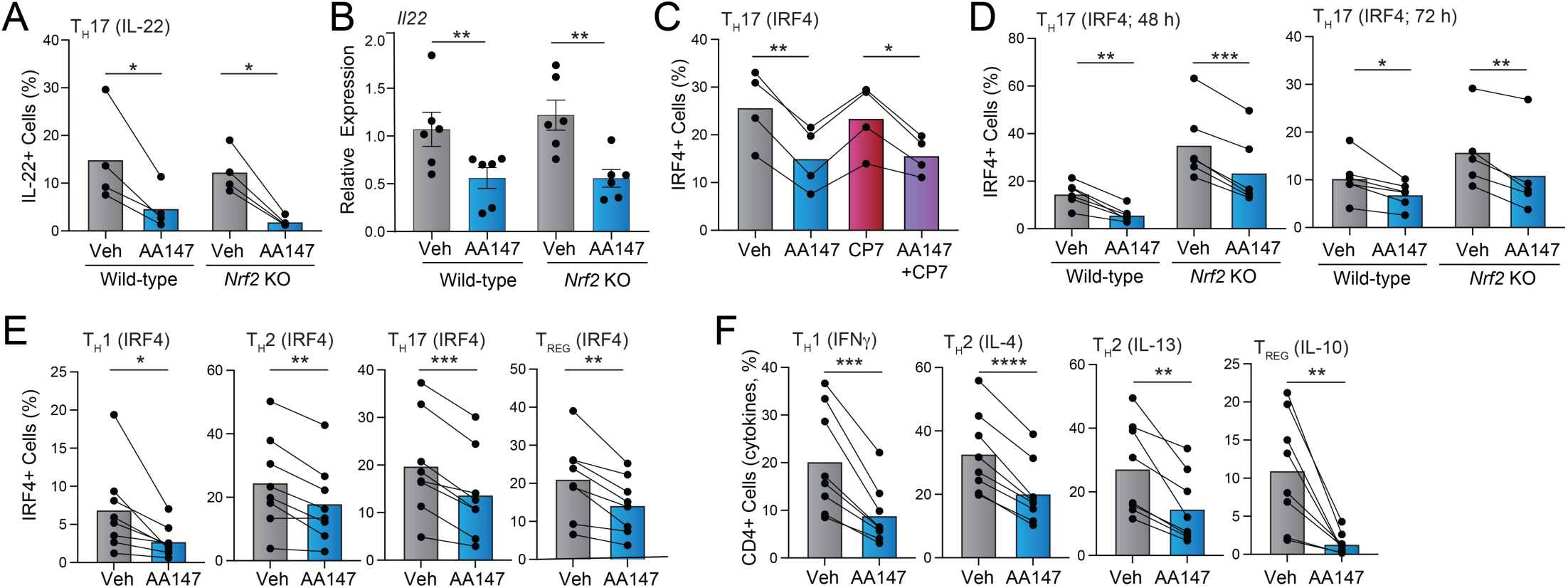
AA147 reduces IRF4 in CD4^+^ T cells through an NRF2-independent mechanism. **A.** Percentage of CD4^+^ IL-22^+^ in T_H_17 cells differentiated from wild-type or *Nrf2* KO mice for 72 h in the presence of AA147 (30 µM). Data from CD4^+^ T cells isolated from n=4 mice is shown. **B**. Expression, measured by RT-qPCR, of *Il22* in CD4^+^ T cells from wild-type or *Nrf2* KO mice differentiated for 72 h in T_H_17 polarising conditions in the presence or absence of AA147 (30 µM). Data from CD4^+^ T cells isolated from n=6 mice is shown. **C**. Percentage of CD4^+^ IRF4^+^ in T_H_17 cells differentiated for 72 h in the presence of AA147 (30 µM) and/or CP7 (3 µM), as indicated. Data from CD4^+^ T cells isolated from n=4 mice is shown. **D**. Percentage of CD4^+^ IRF4^+^ cells differentiated from wild-type or *Nrf2* KO mice for 48 h (left) or 72 h (right) in the presence of AA147 (30 µM). Data from CD4^+^ T cells isolated from n=6 mice is shown. **E**. Percentage of CD4^+^ IRF4+ in cells differentiated for 72 h in T_H_1, T_H_2, T_H_17, or T_REG_ polarising conditions, as indicated, in the presence or absence of AA147 (30 µM). Data from CD4^+^ T cells isolated from n=8 mice is shown. **F**. Percentage of CD4^+^ IFNγ^+^, IL-4^+^, IL-13^+^, or IL-10^+^ cells differentiated for 72 h in T_H_1, T_H_2, or T_REG_ polarising conditions, as indicated, in the presence or absence of AA147 (30 µM). Data from CD4^+^ T cells isolated from n=8 mice is shown. *p<0.05, **p<0.01, ***p<0.005 for one-way ANOVA (panel **C**) or paired t-tests (panels **A, B, D, E, F**) or unpaired t-test (normalised data in panel **A**). Data described in panels A-Fare provided in **Source Data** Figure 5.

A primary function of IRF4 is to control the expression of cytokines across various T cell lineages.(Huber and Lohoff, 2014) While AA147 does not influence expression of lineage-defining transcription factors in other CD4^+^ T cell lineages (**Fig. S1C,D**), we found that treatment with AA147 reduced IRF4 protein levels in CD4^+^ T cells differentiated to T_H_1, T_H_2, or T_REG_ cell lineages (**Fig. 5E**, **Fig. S5D**). We also found that AA147 treatment reduced the expression of effector cytokines from T_H_1 (IFNγ), T_H_2 (IL-4, IL-13), and T_REG_ (IL-10) cells, some of which have previously been linked to IRF4 activity (**Fig. 5F**, **Fig. S5E**).(Lohoff et al., 2002; Mahnke et al., 2016; Rengarajan et al., 2002) Collectively, these results indicate that AA147 treatment broadly reduces IRF4 expression across T cell subsets to decrease expression of effector cytokines across these different lineages.

## DISCUSSION

T_H_17 cells coordinate mucosal barrier defence and neutrophil-mediated immunity against extracellular pathogens and fungi. However, T_H_17 cells have been found to be pathogenic in a variety of autoimmune and pro-inflammatory diseases including psoriasis, inflammatory bowel disease (IBD) and multiple sclerosis (MS).(Xu Lou et al., 2025) The therapeutic promise of targeting this subset of cells has been demonstrated by the clinical success of biologics such as secukinumab and risankizumab, which target IL-17A and IL-23 respectively, to suppress T_H_17 cell activity and development.(Yang et al., 2014) However, a substantial proportion of patients fail to respond adequately to these treatments over time, highlighting the need for complementary therapeutic approaches to mitigate pathogenic T_H_17 cell activity.(Yang et al., 2014)

Here, we show that the metabolically activated proteostasis regulator AA147 selectively suppresses differentiation of T_H_17 cells relative to other CD4^+^ T cell lineages, as defined by the expression of their lineage-defining transcription factors. This reduction in differentiation is partially mediated by the AA147-dependent activation of the NRF2 antioxidant response. This is consistent with previous results highlighting the potential for NRF2 to suppress T_H_17 cell differentiation.(Wu and Zhong, 2024) However, previous studies suggest that NRF2 activation reduced T_H_17 cell differentiation through mechanisms involving impaired activity of the upstream transcription factor pSTAT3 and subsequent reductions in the expression of *Rorc.*(Zhao et al., n.d.) In contrast, we show that AA147-dependent NRF2 activation does not reduce pSTAT3 levels or expression of *Rorc*. Instead, our results support a model in which AA147 reduces intracellular ROS and promotes proteasomal degradation of RORγt, therefore suppressing T_H_17 cell differentiation. These findings demonstrate the potential for pharmacological activators of NRF2 that work through different mechanisms to influence T_H_17 cell differentiation by targeting distinct steps in the differentiation process.

In addition to AA147-dependent reductions in RORγt, we found that AA147 suppresses IRF4 expression across CD4^+^ T subsets through a mechanism independent of both ATF6 and NRF2 signalling. Consistent with the established role of IRF4 in T cell cytokine production, reductions in IRF4 were accompanied by decreased production of specific effector cytokines in all tested T cell subsets including T_H_1, T_H_2, T_H_17, and T_REG_ cells (Lohoff et al., 2002; Rengarajan et al., 2002; Mahnke et al., 2016). However, IRF4 reductions did not correlate with the reduced expression of all effector cytokines, as AA147 treatment did not reduce IL-17A expression in NRF2-deficient CD4^+^ T cells. While the mechanistic basis of AA147-dependent reductions in IRF4 expression remains under active investigation, these results demonstrate an opportunity to more broadly influence CD4^+^ T cell activity across different subsets through AA147-dependent dampening of specific effector cytokines.

The ability for AA147 to activate ATF6 and/or NRF2 signalling has proven highly beneficial for mitigating multiple tissue-specific pathologies in cellular and mouse models of many different diseases. Notably, AA147-dependent activation of ATF6 and NRF2 signalling has been shown to promote the adaptive remodelling of neurons, oligodendrocytes, and microglia in mouse models of ischemic stroke and the EAE mouse model of MS, both conditions in which T_H_17 cell inflammatory responses contribute to disease progression.(Yuan et al., 2022; Aksu et al., 2025) This suggests that apart from the protective remodelling of these neuroglial cell types AA147 could alleviate disease progression by suppressing pro-inflammatory T_H_17 cell activity. Our results highlight a new potential therapeutic opportunity to target pro-inflammatory T_H_17 cell responses that could integrate with AA147-dependent remodelling of other cell types and/or tissues to correct pathologic defects in multi-tissue inflammatory diseases including ischemic stroke and MS.

As we advance this project, we are continuing to probe the mechanistic basis by which AA147 promotes the adaptive remodelling of T cell differentiation and activity. Notably, we are focused on identifying the specific mechanism by which AA147 reduces IRF4 protein levels to suppress the expression of specific cytokines across T cells lineages, revealing new biological pathways that can be pharmacologically targeted to sensitively remodel T cell responses. Further, we are continuing to develop next generation AA147 analogues with improved pharmacokinetics suitable for suppressing pathologic T_H_17 cell differentiation in the context of pro-inflammatory diseases. With the continued development of this class of metabolically activated proteostasis regulators, we will further demonstrate the translational potential for targeting T_H_17 cell differentiation to reshape immune responses in human disease and establish the unique opportunity provided by these compounds to promote both adaptive immune and tissue remodelling that can integrate to mitigate pathologies implicated in the onset and pathogenesis of many complex diseases.

## METHODS

### Mice

C57BL/6 mice were bred in-house by the Rodent Breeding Colony (RBC) at Scripps Research. B6.129X1-*Nfe2l2^tm1Ywk^*/J were purchased from Jackson Laboratory (Strain #:017009). All mice were used at 8- to 12-weeks of age. All animal experiments were performed in accordance with protocols approved by the Institutional Animal Care and Use Committee (IACUC) and complied with Scripps Research institutional guidelines.

### T cell differentiation assays

Plates were coated with Rabbit Anti-Hamster IgG(H+L), Mouse/Rat SP ads-UNLB overnight at 4°C. Naïve CD4^+^ T cells were isolated from splenocytes using MojoSort Mouse Naïve CD4 T Cell Isolation Kit (BioLegend). Isolated T cells were activated with anti-CD3 (145-2C11, 0.25 µg/mL, BioXCell) and anti-CD28 (37.51, 1 µg/mL, BioXCell) and differentiated into T_H_1 cells with rmIL-12 (10 ng/mL, Thermo Fisher), rhIL-2 (50 U/mL, Thermo Fisher) and anti-IL-4 (11B11, 2 µg/mL, BioXCell). T_H_2 cells were differentiated with rhIL-2 (50 U/mL, Thermo Fisher), rmIL-4 (2 ng/mL, Thermo Fisher), and anti-IFNγ (XMG1.2, 2 µg/mL, BioXCell). T_H_17 cells were differentiated with anti-IL-4 (2 µg/mL, BioXCell), and anti-IFNγ (2 µg/mL, BioXCell) with either rhTGFβ (0.3 ng/mL, Thermo Fisher) and rmIL-6 (10 ng/mL, Thermo Fisher) or IL-1β (20 ng/mL, Thermo Fisher), IL-23 (25 ng/mL, Thermo Fisher. T_REG_ cells were differentiated with rhIL-2 (50 U/mL, Thermo Fisher), rhTGFβ (5 ng/mL, Thermo Fisher), anti-IL-4 (2 µg/mL, BioXCell), and anti-IFNγ (2 µg/mL, BioXCell) for 3 days, as previously described. Cells were cultured in complete RPMI 1640 medium with L-glutamine supplemented with 10 mM HEPES, 2 mM L-glutamine, 1 mM sodium pyruvate, 100 U/mL penicillin, 100 ug/mL streptomycin, 50 ug/mL gentamicin and 10% heat inactivated fetal bovine serum (FBS) for the initial activation step. Cells were plated with compounds at doses indicated. 24 h after initial T cell activation, 20 µM beta-mercaptoethanol (BME) was supplemented. Cells were harvested at the time point indicated for either mRNA, transcription factor or cytokine analysis to assess T cell differentiation.

### Compounds

AA147^yne^ was synthesized and characterised as previously described. (Kline et al., 2023) Other compounds were purchased from the following vendors: AA147 (MedChem Express; HY-124293), desmosterol (Sigma; D6513), Ceapin-A7 (Sigma; SML2330), ISRIB (Sigma; SML0843), KI696 (a kind gift from Michael Bollong, Scripps), and bortezomib (Fisher, 50-431-40001). All compounds were dissolved in water or DMSO and used as indicated.

### Flow Cytometry staining

For cytokine analysis, cells were harvested and rested for 2 hours, following which cells were stimulated with PMA (50 ng/mL) and ionomycin (1 µg/mL) in the presence of brefeldin A (1 µg/mL) and monensin (1ug/mL) to promote intracellular cytokine retention for 3 h. Intracellular staining was done using labelled antibodies after fixation and permeabilisation according to manufacturer’s instructions (CytoFix/CytoPerm kit, BD Biosciences). For transcription factor analysis, intracellular staining was done using labelled antibodies after fixation and permeabilisation according to manufacturer’s instructions (Foxp3 transcription factor kit, ThermoFisher). Data was obtained using a Cytek Aurora spectral flow cytometer and analyzed by FloJo v. 10 software. The following antibodies were used: anti-CD4 (BUV496, GK1.5, #612952, BD), anti-CD90 (BUV563, 53-2.1, #741213, BD), anti-CD8a (BUV737, 53-6.7, #612759, BD), anti-CD45 (BV570, 30-F11, #103136, Biolegend), anti-CD44 (BV650, IM7, #103049, Biolegend), anti-IL-17A (efluor450, eBio17B7, #48-7177-82, ThermoFisher), anti-IFNγ (BV711, XMG1.2, #505836, Biolegend), anti-IL-13 (PE-efluor610, eBio13A, #61-7133-82, ThermoFisher), anti-IL-4 (PE, 11B11, #12-7041-82, ThermoFisher), anti-IL-22 (PerCP-efluor710, 1H8PWSR, #46-7221-80, ThermoFisher), anti-IL-10 (PE-Cy7, JES5-16E3, #505026, Biolegend), anti-T-bet (FITC, 4B10, #644812, Biolegend), anti-GATA3 (PerCP-efluor 710, TWAJ, #46-9966-41, ThermoFisher), anti-Foxp3 (PE-Cy5.5, FJK-16s, #35-5773-82, ThermoFisher) and anti-RORγt (PE-Cy7, Q31-378, #567305, BD).

### DCFDA staining

Cells were incubated with 10uM DCFDA for 30mins at 37C, followed by staining with Sytox Blue Dye prior to analysis using a Cytek Aurora spectral flow cytometer and analyzed by FlowJo v. 10 software. A FMO control was used to assist with gating and the mean fluorescent intensity was used to quantify changes.

### Immunoblotting

Cells were lysed in ice-cold RIPA buffer with protease and phosphatase inhibitors and benzonase. Protein concentrations were determined by DC Protein Assay (BioRad). Normalised protein concentrations were run on 10% SDS-PAGE gels. Proteins were transferred to nitrocellulose membranes using a wet transfer system, followed by blocking in 5% (w/v) non-fat dry milk in Tris-buffered saline containing 0.1% Tween-20 (TBST). Membranes were incubated with primary antibodies diluted in BSA-azide overnight at 4 °C, following which they were incubated with species-appropriate secondary antibodies for 1 h at room temperature. Membranes were imaged and quantified on Licor. Primary antibodies used: Mouse anti-STAT3 (Cell Signalling, #9139), Rabbit anti-Phospho-STAT3 (Tyr705) (Cell Signalling, #9145), Rabbit anti-β-Actin (Cell Signalling, #8457). Secondary antibodies used: IRDye 800CW Goat Anti-Rabbit (LICOR Biotech, #926-32211), IRDye 680CW Goat Anti-Mouse (LICOR Biotech, #NC0252290)

### Protein labelling with rhodamine azide

Cells were cultured in complete RPMI medium supplemented with 10% fetal bovine serum (FBS). Cells were treated with AA147 for 6 h, following which they were treated with AA147yne for 18 h. Cells were harvested in ice-cold PBS and lysed in RIPA buffer with protease and phosphatase inhibitors and benzonase. Click chemistry reactions were performed by incubating clarified lysates with a freshly prepared master mix containing tris(benzyltriazolylmethyl)amine (TBTA; Click Chemistry Tools, 1061-100), tris(2-carboxyethyl)phosphine (TCEP; Sigma, 75259), rhodamine-azide (Vector Labs), and CuSO₄. Reactions were carried out at room temperature for 1 h with intermittent vortexing every 15 min. 4X SDS loading buffer was added to the sample and run on an SDS-PAGE gel. A Biorad gel imager was used to image the gel.

### RNA extraction and quantitative real-time reverse transcription PCR

Total RNA was extracted from pelleted cells using Zymo Research Quick-RNA Miniprep Kit according to manufacturer’s instructions. Complementary DNA (cDNA) was generated using High-Capacity cDNA Reverse Transcription Kit (Applied Biosystems). Quantitative real-time reverse transcription PCR was performed on a QuantStudio 5 Real-Time PCR system (Thermo Fisher) using SYBR Green technology. Results were analyzed and presented as the fold reduction to the internal control primer for the housekeeping gene, *Actb.* PCR primers (5’-3’) for mouse gene sequences were as follows:

*Actb*-F: CATTGCTGACAGGATGCAGAAGG
*Actb*-R: TGCTGGAAGGTGGACAGTGAGG
*Rorc-*F: ACCTCCACTGCCAGCTGTGTGCTGTC
*Rorc-*R: TCATTTCTGCACTTCTGCATGTAGACTGTCCC
*Il17a-*F: CAGACTACCTCAACCGTTCCAC
*Il17a-*R: TCCAGCTTTCCCTCCGCATTGA
*Il17f-*F: AACCAGGGCATTTCTGTCCCAC
*Il17f-*R: GGCATTGATGCAGCCTGAGTGT
*Il22-*F: GCTTGAGGTGTCCAACTTCCAG
*Il22-*R: ACTCCTCGGAACAGTTTCTCCC
*Hspa5-*F: GTCCAGGCTGGTGTCCTCTC
*Hspa5*-R: GATTATCGGAAGCCGTGGAG

### RNA-Sequencing (RNAseq)

Purified RNA (50 ng/µL) was prepared using Zymo Research Quick-RNA Miniprep Kit according to manufacturer’s instructions and submitted to Plasmidsaurus for sequencing and analysis. To assess expression of gene sets regulated by stress-responsive signalling pathways, we used a previously published gene set profiling approach. (Grandjean et al., 2019) A *Rorc* target gene list was compiled from previous literature.(Xiao et al., 2014b) Values were Z-normalised across samples prior to comparison, allowing relative changes to be assessed and compared. The complete RNAseq data is deposited in gene expression omnibus (GEO) as GSE344840.

### TMT quantitative proteomics

Cell pellets from T_H_17 cells differentiated in the presence or absence of AA147 were submitted to the mass spectrometry facility. Samples were Tandem Mass Tagged (TMT) and run on the mass spectrometer. Normalised peptide/protein intensity values across six channels (three vehicle treated and three AA147 treated replications) were collected. Normalised intensities were averaged within each condition on a per protein basis, from which fold change was derived as the mean AA147 value divided by the mean vehicle value. Group differences were treated using paired t-tests. Functional annotation was performed via the DAVID Bioinformatics Database to produce Gene Ontology (GO) terms and KEGG pathways. The complete TMT proteomic data is deposited in MassIVE as MSV000103013.

## Supporting information

Source Data File for Figure 1

Source Data File for Figure 2

Source Data File for Figure 3

Source Data File for Figure 4

Source Data File for Figure 5

Source Data File for Figure S1

Source Data File for Figure S2

Source Data File for Figure S3

Source Data File for Figure S4

Source Data File for Figure S5

## ACKNOWLEDGEMENTS

We would like to thank Antonio Pinto and Jolene Diedrich in the Scripps Multi-Omics Core for help with the proteomic analysis. We thank the National Institutes of Health (AG046495 to RLW and GM146865 to MB) and the Prebys Foundation (to AM) for funding. PC was funded by the Ellen Brown Scripps Foundation and the Skaggs-Oxford programme. S.D. was supported by an Irvington Postdoctoral Fellowship from the Cancer Research Institute.

---

**Figure S1.**
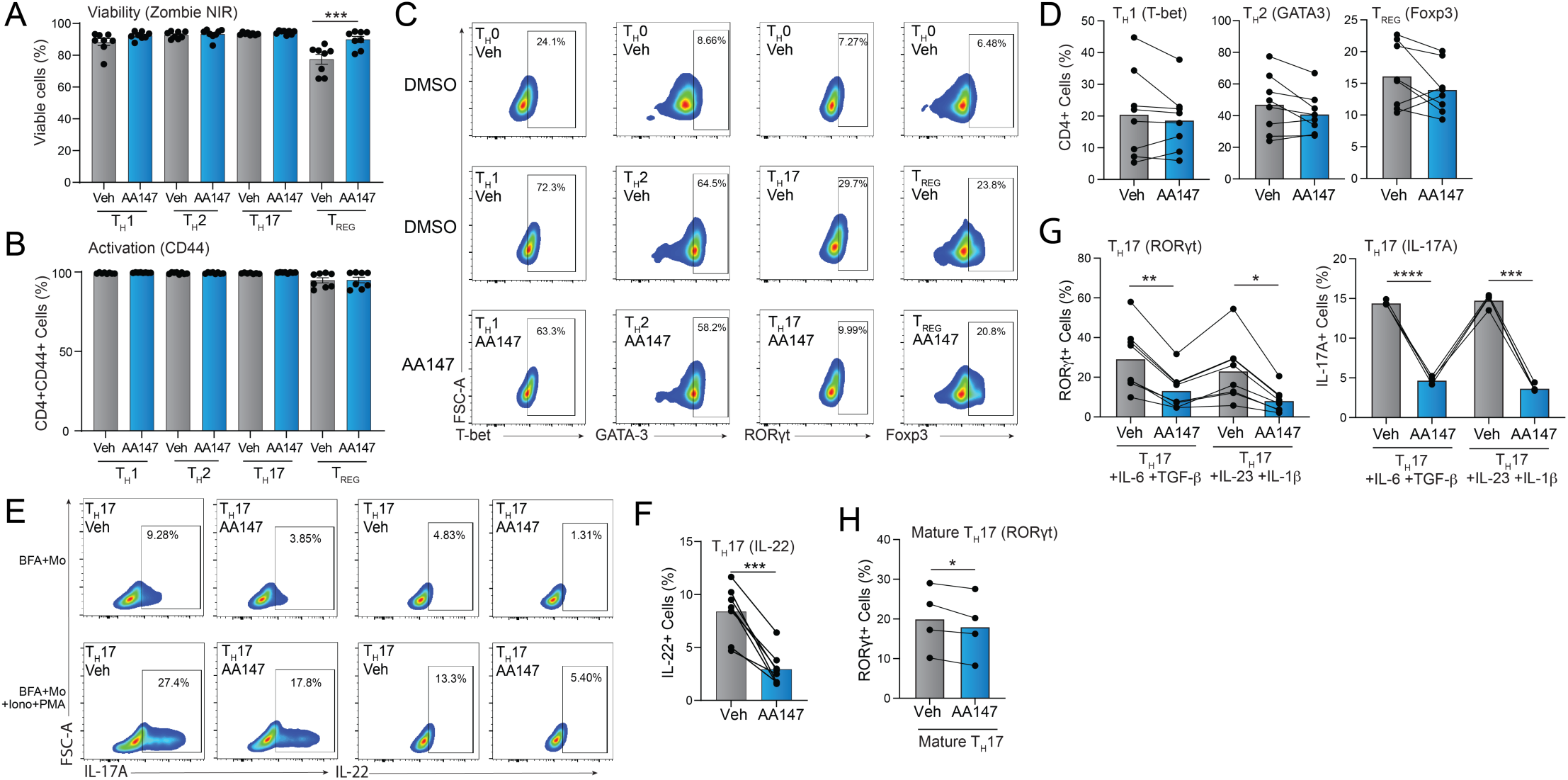
(Supplement to Figure 1). AA147 preferentially decreases differentiation of CD4^+^ T_H_17 cells. **A,B.** Viability, measured by Zombie NIR, (**A**), and activation, measured by CD44 staining (**B**), of CD4^+^ T cells differentiated for 72 h in T_H_1, T_H_2, T_H_17 or T_REG_ polarising conditions in the absence or presence of AA147 (30 µM). **C.** Representative flow cytometry images of the indicated lineage-defining transcription factors in activated CD4^+^ T cells differentiated for 72 h in T_H_1 (T-bet), T_H_2 (GATA3), T_H_17 (RORgt), or T_REG_ (Foxp3) polarising conditions in the presence or absence of AA147. **D**. Percentage of CD4^+^ T-bet^+^ (T_H_1, left), CD4^+^ GATA3^+^ (T_H_2, middle), or CD4^+^ Foxp3^+^ (T_REG,_ right) T cells differentiated in the presence of vehicle or AA147 (30 µM) for 72 h. Data are shown for CD4^+^ naïve T cells isolated from n=8 mice. **E.** Representative flow cytometry images for CD4^+^ IL-17A^+^ (left) and CD4^+^ IL-22^+^ (right) T cells differentiated for 72 h in T_H_17 polarising conditions and then rested for 2 h followed by re-stimulation for 3 h by treatment with phorbol 12-myristate 13-acetate (PMA, 50 ng/mL), ionomycin (Iono, 1 µg/mL), brefeldin A (BFA, 1 µg/mL), and monensin (Mo, 1 µg/mL). Unstimulated cells were rested for 2 h followed by treatment for 3 h with BFA (1 µg/mL) and Mo (1 µg/mL) are shown as a control. **F**. Percentage of CD4^+^ IL-22^+^ T cells differentiated in the presence of vehicle or AA147 (30 µM) for 72 h. Data are shown for CD4^+^ naïve T cells isolated from n=8 mice. **G**. Percentage of CD4^+^ RORgt^+^ (left) or CD4^+^ IL-17A^+^ (right) T cells differentiated for 72 h in the indicated T_H_17 polarising cytokine combination in the absence or presence of AA147 (30 µM). Data are shown for CD4^+^ naïve T cells isolated from n=8 mice. **H**. Percentage of CD4^+^ RORgt^+^ T cells differentiated for 72 h in T_H_17 polarising conditions and then treated for an additional 72 h with AA147 (30 µM). Data are shown for CD4^+^ T cells isolated from n=4 mice. *p<0.05, **p<0.01, ***p<0.005, ****p<0.001 for one-way ANOVA (panels **A**, **G**) or paired t-test (panels **D, F**, **H**). Data described in panels A, B, D, F, G, and H are provided in **Source Data Figure S1.**

**Figure S2.**
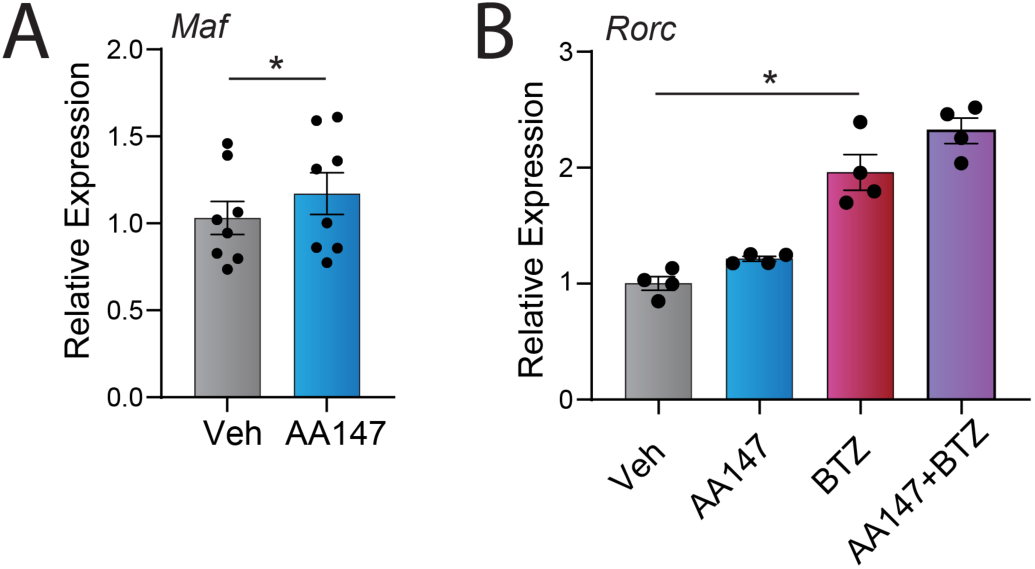
(Supplement to Figure 2). AA147 promotes proteasomal degradation of RORγt. **A.** Expression, measured by RT-qPCR, of the pSTAT3 target gene *Maf* in CD4^+^ T cells differentiated for 72 h in T_H_17 polarising conditions in the presence or absence of AA147 (30 µM). Data are shown for CD4^+^ T cells isolated from n=8 mice. **B**. Expression, measured by RT-qPCR, of *Rorc* in CD4^+^ T cells differentiated for 72 h in T_H_17 polarising conditions in the presence or absence of AA147 (30 µM), followed by treatment with vehicle or BTZ (10 µM) for 4 h, as indicated. Data are shown for CD4^+^ T cells isolated from n=4 mice. *p<0.05 for paired t-test (panel **A**) or one-way ANOVA (panel **B**). Data described in panels A and B are provided in **Source Data Figure S2.**

**Figure S3.**
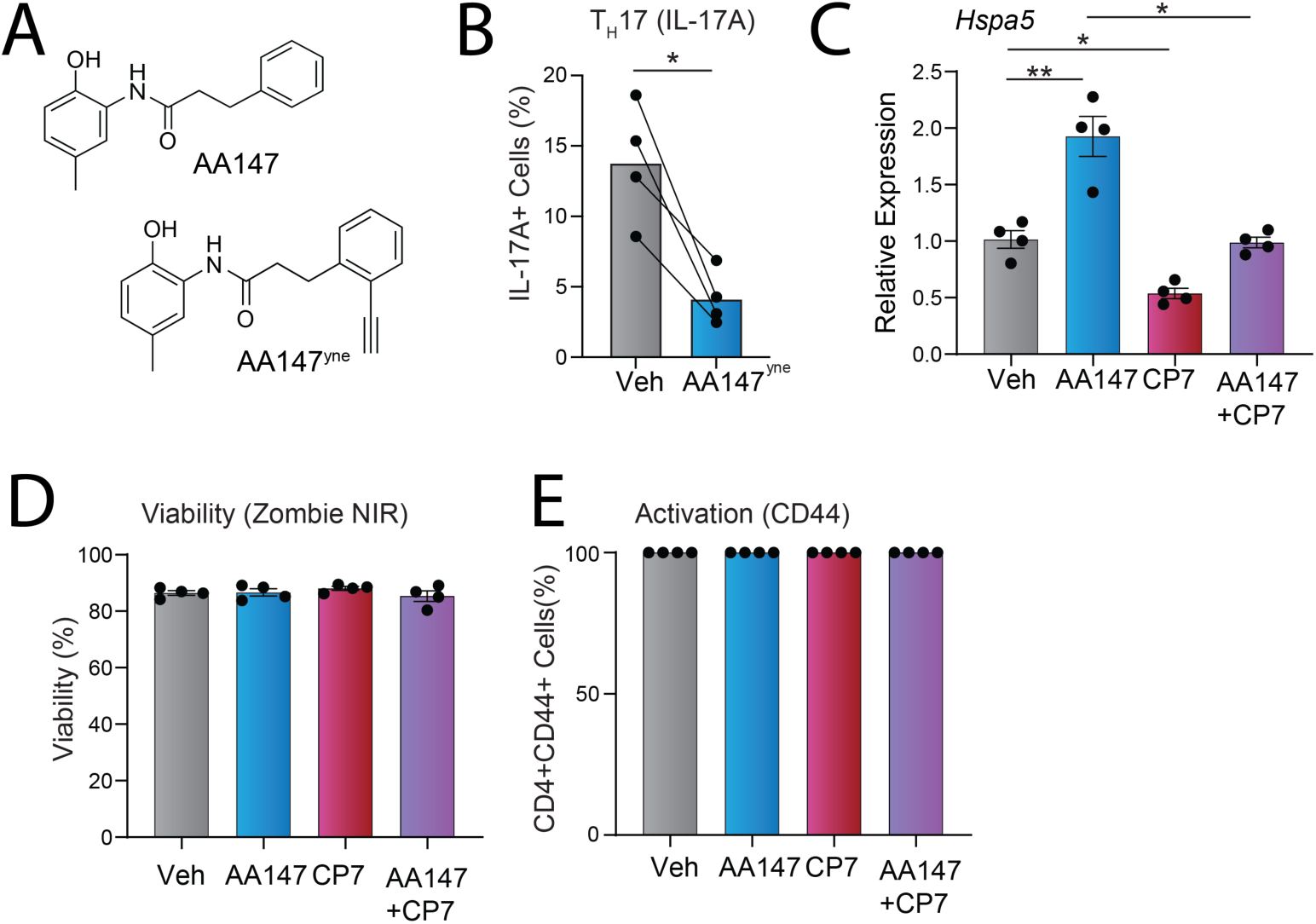
(Supplement to Figure 3). AA147-dependent reductions in T_H_17 differentiation are independent of ATF6 activation. **A.** Structure of AA147 and the alkyne functionalised probe AA147^yne^. **B.** Percentage of CD4^+^ IL-17A^+^ T_H_17 cells differentiated for 72 h in the presence of AA147^yne^ (30 µM). Data are shown for CD4+ T cells isolated from n=4 different mice. **C.** Expression, measured by RT-qPCR, of *Hspa5* in CD4^+^ T cells differentiated for 72 h in T_H_17 polarising conditions in the presence or absence of AA147 (30 µM) and/or CP7 (3 µM). **D, E.** Viability, measured by Zombie NIR (**D**), and activation, measured by CD44 staining (**E**), of CD4^+^ T_H_17 cells differentiated for 72 h in the absence or presence of AA147 (30 µM) and/or CP7 (3 µM), as indicated. Data are shown for CD4^+^ T cells isolated from 4 different mice. *p<0.05, **p<0.01, for one-way ANOVA (panel **C, D, E**) or paired t-test (panel **B**). Data described in panels B-E are provided in **Source Data Figure S3.**

**Figure S4.**
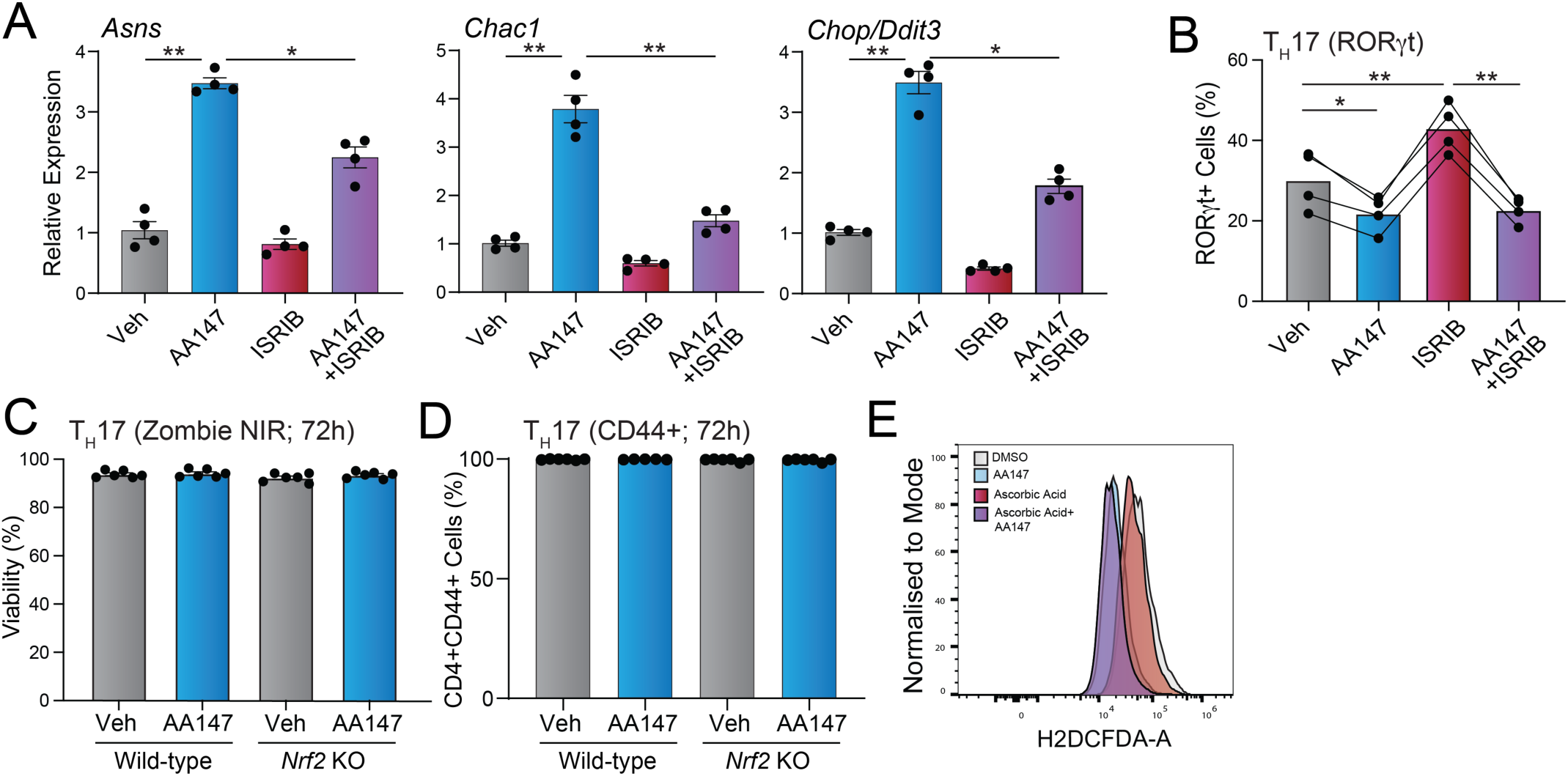
(Supplement to Figure 4). AA147 reduces T_H_17 differentiation through an NRF2-dependent mechanism. **A.** Expression, measured by RT-qPCR, of the ISR target genes *Asns* (left)*, Chac1* (middle) or *Chop/Ddit3* (right) in CD4^+^ T cells differentiated for 72 h in T_H_17 polarising conditions in the presence or absence of AA147 (30 µM) and/or ISRIB (250 nM). Data are shown for CD4^+^ T cells isolated from n=4 mice. **B.** Percentage of CD4^+^ RORgt^+^ T_H_17 cells differentiated for 72 h in the presence or absence of AA147 (30 µM) and/or ISRIB (250 nM). Data for CD4^+^ T cells isolated from n=4 mice are shown. **C, D.** Viability, measured by Zombie NIR, (**C**), and activation, measured by CD44 staining (**D**), of CD4^+^ T cells differentiated from wild-type or *Nrf2* KO mice for 72 h in the T_H_17 polarising conditions in the absence or presence of AA147 (30 µM). Data for CD4^+^ T cells isolated from n=4 mice are shown. **E.** Representative flow cytometry histograms of H_2_DCFDA fluorescence in T_H_17 cells differentiated for 72 h in the presence or absence of AA147 (30 µM) and/or ascorbic acid (250 µM). *p<0.05, **p<0.01, for one-way ANOVA (panels **A**, **B, C, D**). Data described in panels A-D are provided in **Source Data Figure S4.**

**Figure S5.**
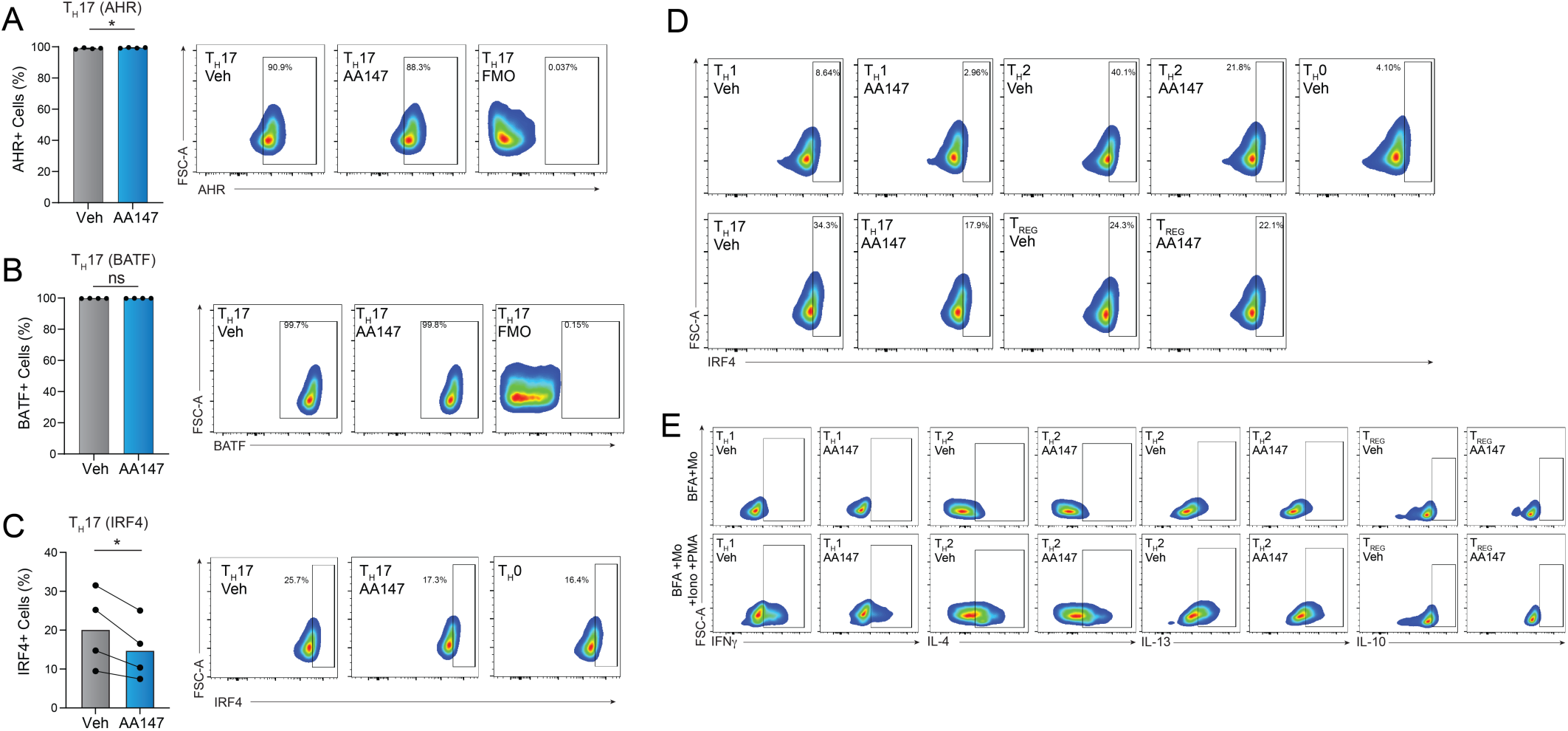
(Supplement to Figure 5). AA147 reduces IRF4 in CD4^+^ T cells through an NRF2-independent mechanism. **A-C.** Representative flow cytometry plot and quantification of CD4^+^ AHR^+^, CD4^+^ BATF^+^, or CD4^+^ IRF4^+^ cells differentiated for 72 h in T_H_17 polarising conditions in the presence or absence of AA147. **D**. Representative flow plots of CD4^+^ IRF4+ in cells differentiated for 72 h in T_H_1, T_H_2, T_H_17, or T_REG_ polarising conditions, as indicated, in the presence or absence of AA147 (30 µM). **E.** Representative flow plots of CD4^+^ IFNγ^+^, IL-4^+^, IL-13^+^, or IL-10^+^ cells differentiated for 72 h in T_H_1, T_H_2, or T_REG_ polarising conditions, as indicated, in the presence or absence of AA147 (30 µM). ***p<0.005, ****p<0.001 for paired t-test (panels **A, B, C**). Data described in panels A-C are provided in **Source Data Figure S5.**

## SUPPLEMENTAL TABLE LEGENDS

**Table S1:** TMT-MS proteomic data from T_H_17 cells differentiated in the presence or absence of AA147 (30 µM) for 72 h. Data shown as relative fold changes with corresponding p values. A p value cut off of 0.05 was used.

**Table S2:** RNAseq data from TH17 cells differentiated in the presence or absence of AA147 (30 µM) for 72 h. A p value cut off of 0.05 was used.

**Table S3:** RNAseq data from TH17 cells differentiated in the presence or absence of AA147 (30 µM) for 72 h showing genes used in the *Rorc* gene list as well as the stress pathway profiling gene list.

## SOURCE DATA FILES

**Source Data Figure 1.** Source data used to generate **Figure 1B, D, E**.

**Source Data Figure S1.** Source data used to generate **Figure S1A, B, D, F, F, G, H.**

**Source Data Figure 2.** Source data used to generate **Figure 2B-E**.

**Source Data Figure S2.** Source data used to generate **Figure S2A, B.**

**Source Data Figure 3.** Source data used to generate **Figure 3C-F**.

**Source Data Figure S3.** Source data used to generate **Figure S3B-E.**

**Source Data Figure 4.** Source data used to generate **Figure 4C-I**.

**Source Data Figure S4.** Source data used to generate **Figure S4-D.**

**Source Data Figure 5.** Source data used to generate **Figure 5A-F**.

**Source Data Figure S5.** Source data used to generate **Figure S5A-C.**

## REFERENCES

Acosta-Alvear, D., Harnoss, J.M., Walter, P., Ashkenazi, A., 2025. Homeostasis control in health and disease by the unfolded protein response. Nat Rev Mol Cell Biol 26, 193–212. 10.1038/s41580-024-00794-0

Adachi, Y., Yamamoto, K., Okada, T., Yoshida, H., Harada, A., Mori, K., 2008. ATF6 is a transcription factor specializing in the regulation of quality control proteins in the endoplasmic reticulum. Cell Struct Funct 33, 75–89. 10.1247/csf.07044

Aksu, M., Kaschke, K., Podojil, J.R., Chiang, M., Steckler, I., Bruce, K., Cogswell, A.C., Schulz, G., Kelly, J.W., Wiseman, R.L., Miller, S.D., Popko, B., Chen, Y., 2025. AA147 Alleviates Symptoms in a Mouse Model of Multiple Sclerosis by Reducing Oligodendrocyte Loss. Glia 73, 1241–1257. 10.1002/glia.70001

Angelini, G., Gardella, S., Ardy, M., Ciriolo, M.R., Filomeni, G., Di Trapani, G., Clarke, F., Sitia, R., Rubartelli, A., 2002. Antigen-presenting dendritic cells provide the reducing extracellular microenvironment required for T lymphocyte activation. Proceedings of the National Academy of Sciences 99, 1491–1496. 10.1073/pnas.022630299

Blackwood, E.A., Azizi, K., Thuerauf, D.J., Paxman, R.J., Plate, L., Kelly, J.W., Wiseman, R.L., Glembotski, C.C., 2019. Pharmacologic ATF6 activation confers global protection in widespread disease models by reprograming cellular proteostasis. Nat Commun 10, 187. 10.1038/s41467-018-08129-2

Carlson, T.J., Pellerin, A., Djuretic, I.M., Trivigno, C., Koralov, S.B., Rao, A., Sundrud, M.S., 2014. Halofuginone-induced amino acid starvation regulates Stat3-dependent Th17 effector function and reduces established autoimmune inflammation. J Immunol 192, 2167–2176. 10.4049/jimmunol.1302316

Chandwaskar, R., Dalal, R., Gupta, S., Sharma, A., Parashar, D., Kashyap, V.K., Sohal, J.S., Tripathi, S.K., 2024. Dysregulation of T cell response in the pathogenesis of inflammatory bowel disease. Scandinavian Journal of Immunology 100, e13412. 10.1111/sji.13412

Chen, S., Wang, Q., Wang, H., Xia, S., 2023. Endoplasmic reticulum stress in T cell-mediated diseases. Scand J Immunol 98, e13307. 10.1111/sji.13307

Cheng, Q., Choi, H.-J., Wu, Y., Yuan, X., Pugel, A., Tian, L., Hendrix, M., Fu, D., Alimohammadi, R., Liu, C., Yu, X.-Z., 2025. ER stress sensor PERK promotes T cell pathogenicity in GVHD by regulating ER-associated degradation. J Clin Invest 135, e190958. 10.1172/JCI190958

Ciofani, M., Madar, A., Galan, C., Sellars, M., Mace, K., Pauli, F., Agarwal, A., Huang, W., Parkhurst, C.N., Muratet, M., Newberry, K.M., Meadows, S., Greenfield, A., Yang, Y., Jain, P., Kirigin, F.K., Birchmeier, C., Wagner, E.F., Murphy, K.M., Myers, R.M., Bonneau, R., Littman, D.R., 2012. A validated regulatory network for Th17 cell specification. Cell 151, 289–303. 10.1016/j.cell.2012.09.016

Durant, L., Watford, W.T., Ramos, H.L., Laurence, A., Vahedi, G., Wei, L., Takahashi, H., Sun, H.-W., Kanno, Y., Powrie, F., O’Shea, J.J., 2010. Diverse Targets of the Transcription Factor STAT3 Contribute to T Cell Pathogenicity and Homeostasis. Immunity 32, 605–615. 10.1016/j.immuni.2010.05.003

Espinosa, J.R., Wheaton, J.D., Ciofani, M., 2020. In Vitro Differentiation of CD4+ T Cell Effector and Regulatory Subsets, in: Liu, C. (Ed.), T-Cell Receptor Signaling: Methods and Protocols. Springer US, New York, NY, pp. 79–89. 10.1007/978-1-0716-0266-9_7

Farchione, A.J., Cheon, H., Vremec, D., Neumann, J., Verstappen, G.M., Hardy, M.Y., Margetts, M.B., Howson, L.J., Henneken, L.M., Forde, M., Tye-Din, J.A., Heinzel, S., Hodgkin, P.D., Bryant, V.L., 2026. Functional immune profiling reveals CD4+ T cell dysregulation in coeliac disease. Immunology & Cell Biology 104, 613–630. 10.1111/imcb.70132

Fasching, P., Stradner, M., Graninger, W., Dejaco, C., Fessler, J., 2017. Therapeutic Potential of Targeting the Th17/Treg Axis in Autoimmune Disorders. Molecules 22, 134. 10.3390/molecules22010134

Gallagher, C.M., Garri, C., Cain, E.L., Ang, K.K.-H., Wilson, C.G., Chen, S., Hearn, B.R., Jaishankar, P., Aranda-Diaz, A., Arkin, M.R., Renslo, A.R., Walter, P., 2016. Ceapins are a new class of unfolded protein response inhibitors, selectively targeting the ATF6α branch. Elife 5, e11878. 10.7554/eLife.11878

Grandjean, J.M.D., Plate, L., Morimoto, R.I., Bollong, M.J., Powers, E.T., Wiseman, R.L., 2019. Deconvoluting Stress-Responsive Proteostasis Signaling Pathways for Pharmacologic Activation Using Targeted RNA Sequencing. ACS Chem. Biol. 14, 784–795. 10.1021/acschembio.9b00134

Hu, X., Wang, Y., Hao, L.-Y., Liu, X., Lesch, C.A., Sanchez, B.M., Wendling, J.M., Morgan, R.W., Aicher, T.D., Carter, L.L., Toogood, P.L., Glick, G.D., 2015. Sterol metabolism controls TH17 differentiation by generating endogenous RORγ agonists. Nat Chem Biol 11, 141–147. 10.1038/nchembio.1714

Huber, M., Lohoff, M., 2014. IRF4 at the crossroads of effector T-cell fate decision. European Journal of Immunology 44, 1886–1895. 10.1002/eji.201344279

Kemp, K.L., Lin, Z., Zhao, F., Gao, B., Song, J., Zhang, K., Fang, D., 2013. The Serine-threonine Kinase Inositol-requiring Enzyme 1α (IRE1α) Promotes IL-4 Production in T Helper Cells*. Journal of Biological Chemistry 288, 33272–33282. 10.1074/jbc.M113.493171

Kline, G.M., Madrazo, N., Cole, C.M., Pannikkat, M., Bollong, M.J., Rosarda, J.D., Kelly, J.W., Wiseman, R.L., 2024. Metabolically activated proteostasis regulators that protect against erastin-induced ferroptosis. RSC Chem Biol 5, 866–876. 10.1039/d4cb00027g

Kline, G.M., Paxman, R.J., Lin, C.-Y., Madrazo, N., Yoon, L., Grandjean, J.M.D., Lee, K., Nugroho, K., Powers, E.T., Wiseman, R.L., Kelly, J.W., 2023. Divergent Proteome Reactivity Influences Arm-Selective Activation of the Unfolded Protein Response by Pharmacological Endoplasmic Reticulum Proteostasis Regulators. ACS Chem Biol 18, 1719–1729. 10.1021/acschembio.3c00042

Kroeger, H., Grandjean, J.M.D., Chiang, W.-C.J., Bindels, D.D., Mastey, R., Okalova, J., Nguyen, A., Powers, E.T., Kelly, J.W., Grimsey, N.J., Michaelides, M., Carroll, J., Wiseman, R.L., Lin, J.H., 2021. ATF6 is essential for human cone photoreceptor development. Proceedings of the National Academy of Sciences 118, e2103196118. 10.1073/pnas.2103196118

Lo, S.-C., Li, X., Henzl, M.T., Beamer, L.J., Hannink, M., 2006. Structure of the Keap1:Nrf2 interface provides mechanistic insight into Nrf2 signaling. EMBO J 25, 3605–3617. 10.1038/sj.emboj.7601243

Lohoff, M., Mittrücker, H.-W., Prechtl, S., Bischof, S., Sommer, F., Kock, S., Ferrick, D.A., Duncan, G.S., Gessner, A., Mak, T.W., 2002. Dysregulated T helper cell differentiation in the absence of interferon regulatory factor 4. Proc Natl Acad Sci U S A 99, 11808–11812. 10.1073/pnas.182425099

Louten, J., Boniface, K., de Waal Malefyt, R., 2009. Development and function of TH17 cells in health and disease. J Allergy Clin Immunol 123, 1004–1011. 10.1016/j.jaci.2009.04.003

Luckheeram, R.V., Zhou, R., Verma, A.D., Xia, B., 2012. CD4+T Cells: Differentiation and Functions. Clin Dev Immunol 2012, 925135. 10.1155/2012/925135

Ma, Q., 2013. Role of Nrf2 in Oxidative Stress and Toxicity. Annu Rev Pharmacol Toxicol 53, 401–426. 10.1146/annurev-pharmtox-011112-140320

Mahnke, J., Schumacher, V., Ahrens, S., Käding, N., Feldhoff, L.M., Huber, M., Rupp, J., Raczkowski, F., Mittrücker, H.-W., 2016. Interferon Regulatory Factor 4 controls TH1 cell effector function and metabolism. Sci Rep 6, 35521. 10.1038/srep35521

Masopust, D., Awasthi, A., Bosselut, R., Brooks, D.G., Buggert, M., Chamoto, K., Cui, W., Dong, C., Farber, D.L., Gebhardt, T., Gerlach, C., Goldrath, A., Greenberg, P.D., Hale, J.S., Hayday, A., Homann, D., Iannacone, M., Jameson, S.C., Jenkins, M.K., Joshi, N.S., Kaech, S.M., Kallies, A., Kamphorst, A.O., Kaplan, M.H., Klenerman, P., Künzli, M., Lanzavecchia, A., Lauer, G.M., Lugli, E., Luster, A.D., Mackay, L.K., McElrath, M.J., Mueller, S.N., Ndhlovu, Z., Ndung’u, T., Ohashi, P.S., Oxenius, A., Pantaleo, G., Pepper, M., Picker, L.J., Quarnstrom, C.F., Reyes-Terán, G., Roederer, M., Rosato, P.C., de Oca, G.S.-M., Sallusto, F., Schumacher, T.N., Schwartz, D.M., Shin, E.-C., Soerens, A.G., Thommen, D.S., Vezys, V., Viola, J.P.B., Walker, B.D., Watts, T.H., Weaver, C.T., Wherry, E.J., Xue, H.-H., Youngblood, B., Ahmed, R., 2026. Guidelines for T cell nomenclature. Nat Rev Immunol 26, 298–313. 10.1038/s41577-025-01238-2

Moser, T., Akgün, K., Proschmann, U., Sellner, J., Ziemssen, T., 2020. The role of TH17 cells in multiple sclerosis: Therapeutic implications. Autoimmun Rev 19, 102647. 10.1016/j.autrev.2020.102647

Ngo, V., Duennwald, M.L., 2022. Nrf2 and Oxidative Stress: A General Overview of Mechanisms and Implications in Human Disease. Antioxidants (Basel) 11, 2345. 10.3390/antiox11122345

Paxman, R., Plate, L., Blackwood, E.A., Glembotski, C., Powers, E.T., Wiseman, R.L., Kelly, J.W., 2018. Pharmacologic ATF6 activating compounds are metabolically activated to selectively modify endoplasmic reticulum proteins. eLife 7, e37168. 10.7554/eLife.37168

Pino, S.C., O’Sullivan-Murphy, B., Lidstone, E.A., Thornley, T.B., Jurczyk, A., Urano, F., Greiner, D.L., Mordes, J.P., Rossini, A.A., Bortell, R., 2008. Protein kinase C signaling during T cell activation induces the endoplasmic reticulum stress response. Cell Stress and Chaperones 13, 421–434. 10.1007/s12192-008-0038-0

Plate, L., Cooley, C.B., Chen, J.J., Paxman, R.J., Gallagher, C.M., Madoux, F., Genereux, J.C., Dobbs, W., Garza, D., Spicer, T.P., Scampavia, L., Brown, S.J., Rosen, H., Powers, E.T., Walter, P., Hodder, P., Wiseman, R.L., Kelly, J.W., 2016. Small molecule proteostasis regulators that reprogram the ER to reduce extracellular protein aggregation. eLife 5, e15550. 10.7554/eLife.15550

Read, A., Schröder, M., 2021. The Unfolded Protein Response: An Overview. Biology (Basel) 10, 384. 10.3390/biology10050384

Rengarajan, J., Mowen, K.A., McBride, K.D., Smith, E.D., Singh, H., Glimcher, L.H., 2002. Interferon Regulatory Factor 4 (IRF4) Interacts with NFATc2 to Modulate Interleukin 4 Gene Expression. The Journal of Experimental Medicine 195, 1003–1012. 10.1084/jem.20011128

Rosarda, J.D., Baron, K.R., Nutsch, K., Kline, G.M., Stanton, C., Kelly, J.W., Bollong, M.J., Wiseman, R.L., 2021. Metabolically Activated Proteostasis Regulators Protect against Glutamate Toxicity by Activating NRF2. ACS Chem Biol 16, 2852–2863. 10.1021/acschembio.1c00810

Rosarda, J.D., Giles, S., Harkins-Perry, S., Mills, E.A., Friedlander, M., Wiseman, R.L., Eade, K.T., 2023. Imbalanced unfolded protein response signaling contributes to 1-deoxysphingolipid retinal toxicity. Nat Commun 14, 4119. 10.1038/s41467-023-39775-w

Rutz, S., Eidenschenk, C., Kiefer, J.R., Ouyang, W., 2016. Post-translational regulation of RORγt—A therapeutic target for the modulation of interleukin-17-mediated responses in autoimmune diseases. Cytokine & Growth Factor Reviews, Special Issue:Cytokines 2015 30, 1–17. 10.1016/j.cytogfr.2016.07.004

Saravia, J., Chapman, N.M., Chi, H., 2019. Helper T cell differentiation. Cell Mol Immunol 16, 634–643. 10.1038/s41423-019-0220-6

Schnell, A., Littman, D.R., Kuchroo, V.K., 2023. TH17 cell heterogeneity and its role in tissue inflammation. Nat Immunol 24, 19–29. 10.1038/s41590-022-01387-9

Sekine, Y., Zyryanova, A., Crespillo-Casado, A., Fischer, P.M., Harding, H.P., Ron, D., 2015. Stress responses. Mutations in a translation initiation factor identify the target of a memory-enhancing compound. Science 348, 1027–1030. 10.1126/science.aaa6986

Shoulders, M.D., Ryno, L.M., Genereux, J.C., Moresco, J.J., Tu, P.G., Wu, C., Yates, J.R., Su, A.I., Kelly, J.W., Wiseman, R.L., 2013. Stress-independent activation of XBP1s and/or ATF6 reveals three functionally diverse ER proteostasis environments. Cell Rep 3, 1279–1292. 10.1016/j.celrep.2013.03.024

Shu, P., Liang, H., Zhang, J., Lin, Y., Chen, W., Zhang, D., 2023a. Reactive oxygen species formation and its effect on CD4+ T cell-mediated inflammation. Front Immunol 14, 1199233. 10.3389/fimmu.2023.1199233

Shu, P., Liang, H., Zhang, J., Lin, Y., Chen, W., Zhang, D., 2023b. Reactive oxygen species formation and its effect on CD4+ T cell-mediated inflammation. Front Immunol 14, 1199233. 10.3389/fimmu.2023.1199233

Sidrauski, C., Acosta-Alvear, D., Khoutorsky, A., Vedantham, P., Hearn, B.R., Li, H., Gamache, K., Gallagher, C.M., Ang, K.K.-H., Wilson, C., Okreglak, V., Ashkenazi, A., Hann, B., Nader, K., Arkin, M.R., Renslo, A.R., Sonenberg, N., Walter, P., 2013. Pharmacological brake-release of mRNA translation enhances cognitive memory. Elife 2, e00498. 10.7554/eLife.00498

Sidrauski, C., Tsai, J.C., Kampmann, M., Hearn, B.R., Vedantham, P., Jaishankar, P., Sokabe, M., Mendez, A.S., Newton, B.W., Tang, E.L., Verschueren, E., Johnson, J.R., Krogan, N.J., Fraser, C.S., Weissman, J.S., Renslo, A.R., Walter, P., 2015. Pharmacological dimerization and activation of the exchange factor eIF2B antagonizes the integrated stress response. Elife 4, e07314. 10.7554/eLife.07314

Sligar, C., Sluyter, R., Ooi, L., 2026. Immune imbalance between T helper 1, T helper 17 and regulatory T cells fuels amyotrophic lateral sclerosis pathogenesis: disease trajectory, diagnosis and therapeutic implications. J Neuroinflammation 23, 132. 10.1186/s12974-026-03764-9

Sun, S., Wang, C., Zhao, P., Kline, G.M., Grandjean, J.M.D., Jiang, X., Labaudiniere, R., Wiseman, R.L., Kelly, J.W., Balch, W.E., 2023. Capturing the conversion of the pathogenic alpha-1-antitrypsin fold by ATF6 enhanced proteostasis. Cell Chemical Biology 30, 22–42.e5. 10.1016/j.chembiol.2022.12.004

Sundrud, M.S., Koralov, S.B., Feuerer, M., Calado, D.P., Kozhaya, A.E., Rhule-Smith, A., Lefebvre, R.E., Unutmaz, D., Mazitschek, R., Waldner, H., Whitman, M., Keller, T., Rao, A., 2009. Halofuginone Inhibits TH17 Cell Differentiation by Activating the Amino Acid Starvation Response. Science 324, 1334–1338. 10.1126/science.1172638

Torres, S.E., Gallagher, C.M., Plate, L., Gupta, M., Liem, C.R., Guo, X., Tian, R., Stroud, R.M., Kampmann, M., Weissman, J.S., Walter, P., 2019. Ceapins block the unfolded protein response sensor ATF6α by inducing a neomorphic inter-organelle tether. Elife 8, e46595. 10.7554/eLife.46595

Vlachos, C., Gaitanis, G., Katsanos, K.H., Christodoulou, D.K., Tsianos, E., Bassukas, I.D., 2016. Psoriasis and inflammatory bowel disease: links and risks. Psoriasis (Auckl) 6, 73–92. 10.2147/PTT.S85194

Wang, M., Cotter, E., Wang, Y.-J., Fu, X., Whittsette, A.L., Lynch, J.W., Wiseman, R.L., Kelly, J.W., Keramidas, A., Mu, T.-W., 2022. Pharmacological activation of ATF6 remodels the proteostasis network to rescue pathogenic GABAA receptors. Cell Biosci 12, 48. 10.1186/s13578-022-00783-w

Wu, D., Zhang, X., Zimmerly, K.M., Wang, R., Livingston, A., Iwawaki, T., Kumar, A., Wu, X., Campen, M., Mandell, M.A., Liu, M., Yang, X.O., n.d. Unconventional Activation of IRE1 Enhances Th17 Responses and Promotes Airway Neutrophilia. Am J Respir Cell Mol Biol 71, 169–181. 10.1165/rcmb.2023-0424OC

Wu, D., Zhang, X., Zimmerly, K.M., Wang, R., Wang, C., Hunter, R., Wu, X., Campen, M., Liu, M., Yang, X.O., 2023. Unfolded protein response factor ATF6 augments T helper cell responses and promotes mixed granulocytic airway inflammation. Mucosal Immunol 16, 499–512. 10.1016/j.mucimm.2023.05.007

Wu, H., Zhong, X., 2024. Nrf2 as a Therapy Target for Th17-Dependent Autoimmune Disease, in: The Role of NRF2 Transcription Factor. IntechOpen. 10.5772/intechopen.1005037

Xiao, S., Yosef, N., Yang, J., Wang, Y., Zhou, L., Zhu, C., Wu, C., Baloglu, E., Schmidt, D., Ramesh, R., Lobera, M., Sundrud, M.S., Tsai, P.-Y., Xiang, Z., Wang, J., Xu, Y., Lin, X., Kretschmer, K., Rahl, P.B., Young, R.A., Zhong, Z., Hafler, D.A., Regev, A., Ghosh, S., Marson, A., Kuchroo, V.K., 2014a. Small-Molecule RORγt Antagonists Inhibit T Helper 17 Cell Transcriptional Network by Divergent Mechanisms. Immunity 40, 477–489. 10.1016/j.immuni.2014.04.004

Xiao, S., Yosef, N., Yang, J., Wang, Y., Zhou, L., Zhu, C., Wu, C., Baloglu, E., Schmidt, D., Ramesh, R., Lobera, M., Sundrud, M.S., Tsai, P.-Y., Xiang, Z., Wang, J., Xu, Y., Lin, X., Kretschmer, K., Rahl, P.B., Young, R.A., Zhong, Z., Hafler, D.A., Regev, A., Ghosh, S., Marson, A., Kuchroo, V.K., 2014b. Small-Molecule RORγt Antagonists Inhibit T Helper 17 Cell Transcriptional Network by Divergent Mechanisms. Immunity 40, 477–489. 10.1016/j.immuni.2014.04.004

Xu Lou, I., Zhou, H., Wan, H., 2025. The critical role of Th17 cells and IL-17A in autoimmune and inflammation-associated neurological diseases: mechanisms and therapeutic perspectives. Front. Immunol. 16. 10.3389/fimmu.2025.1656422

Yan, G., Song, R., Zhang, J., Li, Z., Lu, Z., Liu, Z., Zeng, X., Yao, J., 2024. MIF promotes Th17 cell differentiation in rheumatoid arthritis through ATF6 signal pathway. Mol Med 30, 237. 10.1186/s10020-024-01005-4

Yan, Z., Banerjee, R., 2010. Redox Remodeling as an Immunoregulatory Strategy. Biochemistry 49, 1059. 10.1021/bi902022n

Yang, J., Sundrud, M.S., Skepner, J., Yamagata, T., 2014. Targeting Th17 cells in autoimmune diseases. Trends Pharmacol Sci 35, 493–500. 10.1016/j.tips.2014.07.006

Yasuda, D., Toyoshima, K., Kojima, K., Ishida, H., Kaitoh, K., Imamura, R., Kanamitsu, K., Kojima, H., Funakoshi-Tago, M., Osawa, M., Ohe, T., Hirano, T., 2025. Development of Keap1-Nrf2 Protein–Protein Interaction Inhibitor Activating Intracellular Nrf2 Based on the Naphthalene-2-acetamide Scaffold, and its Anti-Inflammatory Effects. ChemMedChem 20, e202500474. 10.1002/cmdc.202500474

Yuan, Z., Lu, L., Lian, Y., Zhao, Y., Tang, T., Xu, S., Yao, Z., Yu, Z., 2022. AA147 ameliorates post-cardiac arrest cerebral ischemia/reperfusion injury through the co-regulation of the ATF6 and Nrf2 signaling pathways. Front Pharmacol 13, 1028002. 10.3389/fphar.2022.1028002

Zhang, D.D., 2025. Thirty years of NRF2: advances and therapeutic challenges. Nat Rev Drug Discov 24, 421–444. 10.1038/s41573-025-01145-0

Zhang, W., Cao, X., 2025. Unfolded protein responses in T cell immunity. Front. Immunol. 15. 10.3389/fimmu.2024.1515715

Zhao, M., Chen, H., Ding, Q., Xu, X., Yu, B., Huang, Z., 2016. Nuclear Factor Erythroid 2-related Factor 2 Deficiency Exacerbates Lupus Nephritis in B6/lpr mice by Regulating Th17 Cell Function. Sci Rep 6, 38619. 10.1038/srep38619

Zhao, Z., Wang, Y., Gao, Y., Ju, Y., Zhao, Y., Wu, Z., Gao, S., Zhang, B., Pang, X., Zhang, Y., Wang, W., n.d. The PRAK-NRF2 axis promotes the differentiation of Th17 cells by mediating the redox homeostasis and glycolysis. Proc Natl Acad Sci U S A 120, e2212613120. 10.1073/pnas.2212613120

Zhu, J., Yamane, H., Paul, W.E., 2010. Differentiation of Effector CD4 T Cell Populations. Annu Rev Immunol 28, 445–489. 10.1146/annurev-immunol-030409-101212

